# Multi-ensemble coordination between the direct and indirect striatal pathways actuates perceptual decisions

**DOI:** 10.1101/2021.11.16.468594

**Authors:** Lele Cui, Shunhang Tang, Kai Zhao, Jingwei Pan, Zhaoran Zhang, Bailu Si, Ning-long Xu

## Abstract

Sensory-guided decision-making is a vital brain function critically depending on the striatum, a key brain structure transforming sensorimotor information into actions. However, how the two opposing striatal pathways work in concert to select actions during decision-making remains controversial. Here, using cell-type specific two-photon imaging and optogenetic perturbations from the posterior dorsal striatum during decision-making behavior in mice, we uncover the population coding and causal mechanisms of the direct- and indirect-pathway spiny projection neurons (dSPNs and iSPNs) in decision-related action selection. Unexpected from prevailing models, we found that both dSPNs and iSPNs contain divergent subpopulations representing competing choices, and exhibit ensemble-level asymmetry: stronger contralateral dominance in dSPNs than in iSPNs. Such multi-ensemble competition/cooperation causally contributes to decision-related action selection, as supported by systematic optogenetic manipulations and verified by computational modeling. Our results unravel a multi-ensemble coordination mechanism in the striatum for action selection during decision-making.

---

As the largest receptive structure in the basal ganglia, the striatum receives topographically organized excitatory input from virtually the entire cortical and thalamic regions, and provides output to downstream basal ganglia nuclei to influence motor output [1–3], and hence plays a pivotal role in decision-making [4–6]. The posterior tail of the dorsal striatum (TS) in particular receives dense excitatory input from sensory cortex and thalamus [1,2,7–9], and therefore represents as a key node in sensorimotor decision-making. The striatum contains two major subtypes of projection neurons, the dSPNs and iSPNs, giving rise to the direct and indirect pathways innervating distinct downstream nuclei. Earlier studies show that the two striatal pathways play opposing roles in movement control, with the direct pathway promoting movements, while the indirect pathway suppressing movements [10–14]. Based on these properties, classical models postulate that the direct pathway should be active and the indirect pathway inactive during actions [4,10–13,15]. However, recent studies reported that both the dSPNs and iSPNs were concurrently active and often show indistinguishable activity patterns during spontaneous movements [16–20], apparently contradicting the classical models. These controversies raise the puzzling question of how the two concurrently active but opposing pathways coordinate to select from competing actions [4,13,15], and urge a direct investigation of the precise roles of cell-type specific striatal circuits in action selection during decision-making.

Here we employed in vivo deep-brain two-photon imaging to record single-cell level activity over a large population of dSPNs and iSPNs during a precisely controlled auditory-guided decision-making task [21, 22]. We found that both dSPNs and iSPNs comprise subpopulations with heterogeneous preferences to competing choices (contralateral or ipsilateral side), representing a multi-ensemble organization. The ensemble activity of the two pathways differ in both response amplitude and temporal synchronization, with stronger contralateral dominance in dSPNs than in iSPNs. Using cell-type specific bidirectional optogenetic manipulations, we found that dSPNs and iSPNs in play opposite causal roles in the auditory decision-making behavior. Concurrent disinhibition of both dSPNs and iSPNs via inhibiting striatal pavalbumin (PV) interneurons biased choice behavior toward the contralateral side, a causal effect corroborates the prediction from the ensemble-level differences between the two pathways. Based on our observed ensemble activity patterns of dSPNs and iSPNs, we built a computational model to demonstrate the plausibility of the causal contributions to decision-making by coordinated striatal ensembles. These findings reveal a multi-ensemble coordination mechanism in the striatum for action selection during decision-making.

## Results

### Two-photon imaging from TS reveals single-cell heterogeneity of both dSPNs and iSPNs

We trained head-fixed mice to perform an auditory-guided decision-making task [21, 22]. Mice were required to classify sound stimuli (pure tones between 5 and 20 kHz) as belonging to high or low frequency categories (**Fig. 1A**; Supplementary movie 1). The categorical choices were quantified as the probability of licking the left or right lick port in response to various tone frequencies (**Fig. 1B**), allowing for psychophysical measurement of perceptual decisions (**Fig. 1C**).

**Figure 1.**
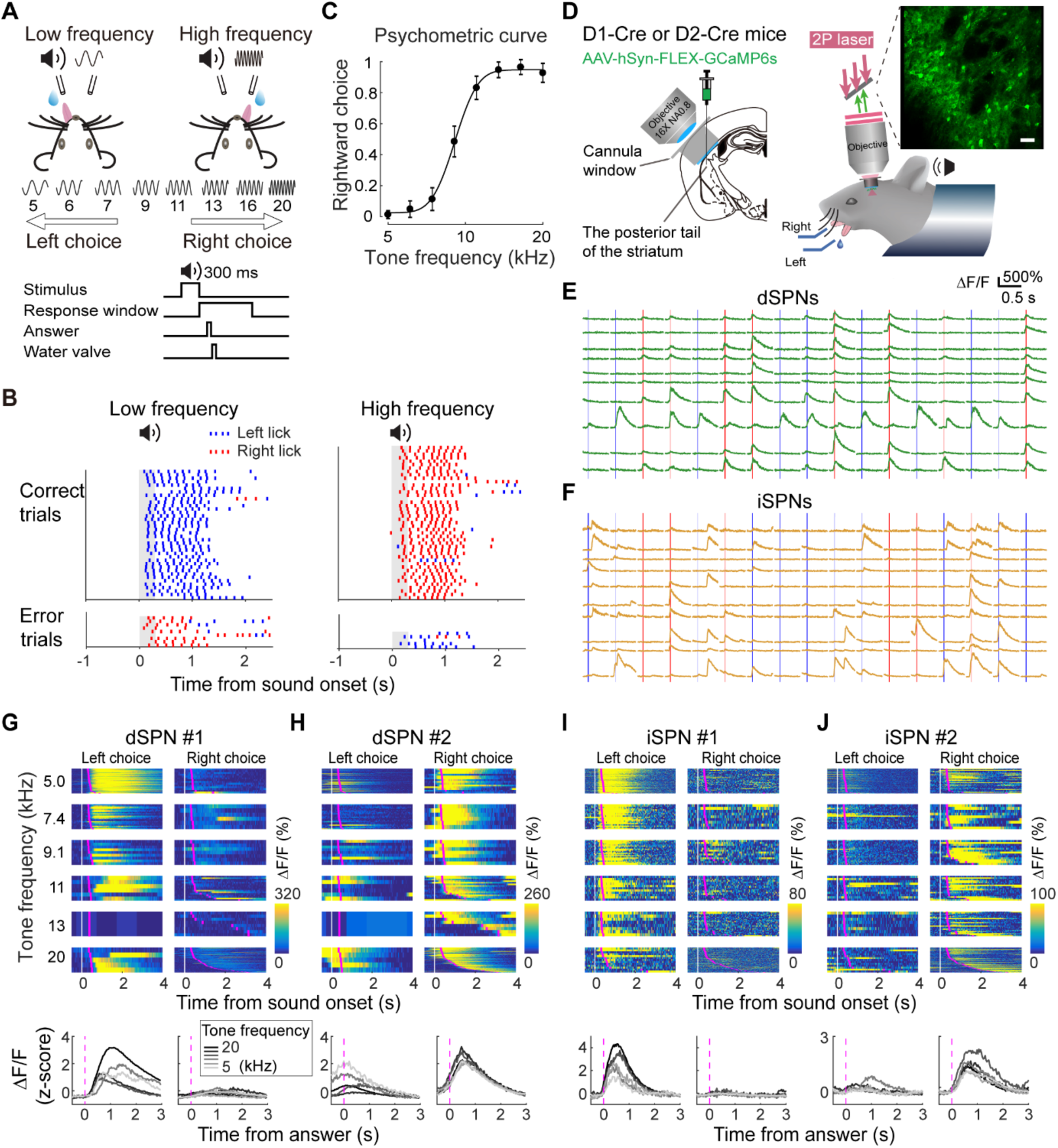
Two-photon imaging from dSPNs and iSPNs during sensorimotor decision task. (A) Top, schematic of the behavioral task. Bottom, temporal structure of the task. (B) Raster plot of lick time from an example session (blue, lick right; red, lick left). Grey shadows indicate the stimulus period. (C) Psychometric function from one example behavioral session. Error bars indicate 95% confidence interval. (D) Schematic showing experimental configuration of *in vivo* two-photon imaging from striatal SPNs labeled with GCaMP6s, through a chronic cannula window in the TS during behavior (see Methods). (E-F) Example traces of calcium signals of dSPNs (E) and iSPNs (F) when mice were performing auditory decision task. The vertical lines indicate mice’s answer time (blue, left choice; red, right choice; light blue and light red, error trials). (G-H) Color raster plot showing calcium signals (ΔF/F) from two example dSPNs with opposite preferences to leftward (G, dSPN #1) and rightward (H, dSPN #2) choice trials in the same imaging field during a behavioral session. Each row in the color raster represents calcium signals from one trial. Blocks of trials were organized according to tone frequency and choice types. White vertical lines, stimulus onset; magenta lines, answer time. Bottom, mean calcium signals averaged from corresponding blocks of trials aligned to answer times. (I-J) Similar as in (G) and (H), for two example iSPNs showing opposite preferences to leftward (I, iSPN #1) and rightward (J, iSPN #2) choices in the same imaging field.

Since striatum receives topographically organized cortical and thalamic inputs [1, 2], to examine the relationship between striatal neurons and auditory decision-making behavior it was important to first identify the subregion that is directly involved in the task. The posterior tail of the striatum (TS) receives input primarily from sensory cortices [1, 2] and was implicated in auditory-related choice behavior [8, 9]. But a direct test of the involvement of TS in auditory decision behavior and its specificity relative to other striatal subregions have been lacking. We therefore first performed optogenetic silencing in both the TS and a dorsal medial region of the striatum (DMS) during task performance. To silence TS, we injected AAV-CAG-ArchT in TS of wild-type mice, and implanted fiber optics above the injection sites (see Methods). We delivered unilateral photostimulation (532 nm, duration, 1 s, starting from sound onset) in the TS in a randomly interleaved subfraction of trials (∼15%), and found that the proportion of contralateral choices was significantly reduced comparing to control trials (Supplementary Fig. 1A-C), indicating an impairment in contralateral action selection. To rule out any non-specific photostimulation effects, we injected AAV-CAG-EGFP to TS in another group of mice and found that the same photostimulation did not produce any significant effect in behavioral performance (Supplementary Fig. 1D). To silence DMS, we injected AAV-CAG-ArchT in a more anterior and medial region of the dorsal striatum (see Methods). The same photostimulation did not significantly change behavioral performance (Supplementary Fig. 1E). These results confirm that TS was specifically required for our auditory decision-making task.

Next we performed *in vivo* two-photon calcium imaging from individual dSPNs or iSPNs through a cannular imaging window in TS (Supplementary Fig. 2; Methods). Genetically encoded calcium sensor GCaMP6s was expressed selectively in dSPNs or iSPNs by injecting AAV-hSyn-FLEX-GCaMP6s in TS of either D1-cre or D2-cre transgenic mice (**Fig. 1D** and Supplementary Fig. 2; see Methods). Task related calcium signals with high signal to noise ratio from individual neurons were reliably detected (**Fig. 1E** and **1F**). We found that activity in individual neurons exhibited clear task modulation, often showing markedly different responses to different choices regardless of stimulus category. Neurons showing preferred responses to either left or right choices (on the contra- or ipsilateral side of the imaged hemisphere) often coexist in simultaneously imaged dSPNs or iSPNs (**Fig. 1G** to **1J**).

We then examined the single-neuron encoding of choice and sensory information across all imaged neurons for both dSPNs and iSPNs by comparing response selectivity in correct and error trials using Receiver Operating Characteristic (ROC) analysis [23]. We found that in both dSPNs and iSPNs, the majority of neurons with significant selectivity are selective to choice rather than to sensory information (Supplementary Fig. 3; Methods), consistent with the notion that striatal projection neurons play important roles in action selection during decision-making. Both dSPNs and iSPNs contain divergent subpopulations showing significant selectivity to either contralateral or ipsilateral choices (Supplementary Fig. 3, Methods), indicating within-cell type heterogeneity in subpopulations of both cell types. Such heterogeneous ensemble organization raises the possibility that previously observed concurrent activation of dSPNs and iSPNs can be due to the activation of different subpopulations of dSPNs and iSPNs preferring the same motor actions, but not necessarily due to uniform co-activation of dSPNs and iSPNs. It hence suggests that the coordination of the direct and indirect pathways can involve more complex competition and cooperation among multiple striatal ensembles from the two hemispheres.

### Ensemble activity shows stronger contralateral dominance in dSPNs than in iSPNs before decision execution

To investigate how different neuronal ensembles in dSPNs and iSPNs coordinate during decision-making, we first analyzed the response amplitude in all SPNs with significant choice selectivity in correct trials prior to animals’ answer time. Overall, there are higher proportions of contralateral preferring neurons (dSPNs, 275/811, 34%; iSPNs, 307/1152, 27%) than ipsilateral preferring neurons (dSPNs, 155/811, 19%; iSPNs, 125/1152, 11%) (**Fig. 2A** and **2B**). By averaging the standard calcium signals across neurons with the same choice preference, we found that the amplitude of responses to preferred choices are stronger in contralateral preferring neurons than in ipsilateral preferring neurons, both in dSPNs (**Fig. 2A**, upper left vs. lower right) and in iSPNs (**Fig. 2B**, upper left vs. lower right). Therefore, the contralateral preferring subpopulations are dominant both in dSPNs and iSPNs, implying that action selection would primarily depend on the competition between contralateral preferring dSPNs and iSPNs. We thus examined the relative strength between these two subpopulations. As shown in **Fig. 2C**, the activation level in dSPNs during contralateral trials is significantly greater than that in iSPNs, beginning from before the answer time (see also **Fig. 2G**). Consistently, the contralateral preference (difference between contra- and ipsilateral responses) is also significantly stronger in dSPNs than in iSPNs, beginning from before the answer time (**Fig. 2D**). In contrast, the difference between ipsilateral preferring dSPNs and iSPNs is relatively minor in their ipsilateral responses (**Fig. 2E** and **2G**) and in their ipsilateral preferences (difference between ipsi- and contralateral responses; **Fig. 2F**).

**Figure 2.**
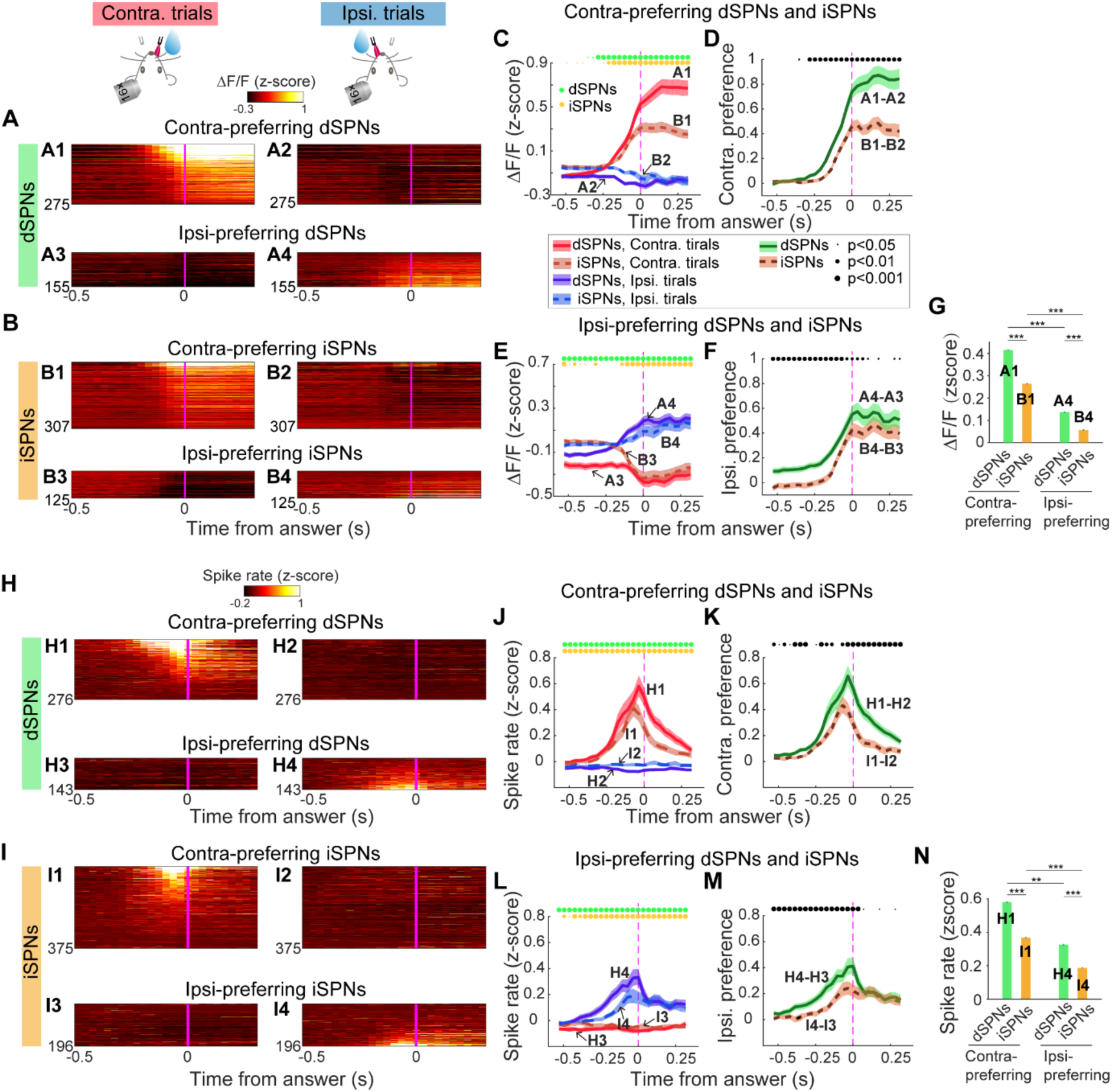
dSPNs and iSPNs comprise subpopulations with preference to contralateral or ipsilateral choices preceding answer time. (A) Color raster plots showing z-scored calcium signals of dSPNs with significant selectivity to contralateral (A1, A2) and ipsilateral choices (A3, A4) before answer time. Each row in the color raster represents mean calcium signals from one neuron averaged from correct contralateral trials (A1, A3) or ipsilateral trials (A2, A4) sorted by SI values. Trials were aligned to answer time (magenta line). (B) Similar as in (A) for iSPNs. (C) Calcium signals averaged across contra-preferring neurons in dSPNs (A1 and A2) and iSPNs (B1 and B2). Shadings indicate bootstrap 95% confidence interval. Statistical difference level across time between contralateral and ipsilateral trials for each subtype of SPNs are indicated as circles above the plots (paired two-sided Wilcoxon signed rank test). (D) Contralateral preference (response difference between contralateral and ipsilateral trials) for contra-preferring dSPNs and iSPNs. Statistical difference level across time between dSPNs and iSPNs are indicated as circles above the plots (two-sided Wilcoxon rank sum test). (E) Similar as in (C), for ipsilateral-preferring dSPNs and iSPNs. (F) Similar as in (D), for ipsilateral preference (response difference between ipsilateral and contralateral trials) in ipsi-preferring dSPNs and iSPNs. (G) Summary of the calcium signals preceding answer time for all choice selective dSPNs and iSPNs in preferred trial types. Shadings indicate bootstrap 95% confidence interval. (H-N) Similar as in (A-F), using inferred spike rate (z-scored) from calcium signal deconvolution.

Taken together, our single-cell resolution population imaging reveals that while multiple subpopulations of SPNs form a complex competition and cooperation system, the dSPNs and iSPNs exhibit structured ensemble-level difference: higher contralateral dominance in dSPNs than in iSPNs. Such differentiation arose before the decision execution time, and is thus likely to contribute to action selection.

To further examine the time course of the response differences between dSPNs and iSPNs, we performed deconvolution of the calcium signals to infer the underlying spike rate changes, which show faster dynamics than original calcium signals (see Methods). We found that the peak of spike rate in response to preferred choices occurred before the answer time in both dSPNs and iSPNs (**Fig. 2H, I, J, L**). The contralateral preferences based on spike rate is also stronger in dSPNs than in iSPNs, with the peaks occurring before the answer time (**Fig. 2K**). This is also the case for ipsilateral preference (**Fig. 2M**). Within dSPNs and iSPNs, the peak spike rate changes in response to preferred choices are also greater in contralateral preferring neurons than in ipsilateral preferring neurons (**Fig. 2N**). Thus, the response differences between dSPNs and iSPNs reached peak level before decision execution, and hence is likely to contribute to decision-related action selection.

### The dSPNs and iSPNs exhibit differential population synchronization

Response strength can be expressed not only in the form of individual neurons’ firing rate, but also in the population-level synchronization between neurons. Two-photon imaging allows us to examine the spatiotemporal activity patterns in simultaneously imaged neurons with single-cell resolution. We asked whether there was a differentiation between dSPNs and iSPNs in their spatiotemporal activity during the task. First, we found that for both dSPNs and iSPNs, nearby neurons showed greater temporal correlation than more distant neurons, with a monotonic dependence on inter-neuronal distances (**Fig. 3A**-**G**). A previous study using single-photon endoscopic imaging during self-paced movements reported similar dependence of the temporal correlation on spatial distance, but the temporal correlations were not directly compared between dSPNs and iSPNs [18]. Here we further compared such inter-neuronal temporal correlations between dSPNs and iSPNs. We found that while both dSPNs and iSPNs showed similar dependence of correlation on inter- neuronal distance, the correlations were overall markedly higher in dSPNs than in iSPNs (**Fig. 3G** and Supplementary Fig. 4). Indeed, two-way repeated-measures ANOVA showed that the inter-neuronal correlations significantly depended on cell types (P<0.01), indicating that the temporal synchronization is higher in simultaneously imaged dSPNs than in iSPNs. This difference cannot be attributed to a higher level of activity in dSPNs, since the overall activity level in dSPN during behavior was not greater but even slightly lower than that in iSPNs (Supplementary Fig. 5).

**Figure 3.**
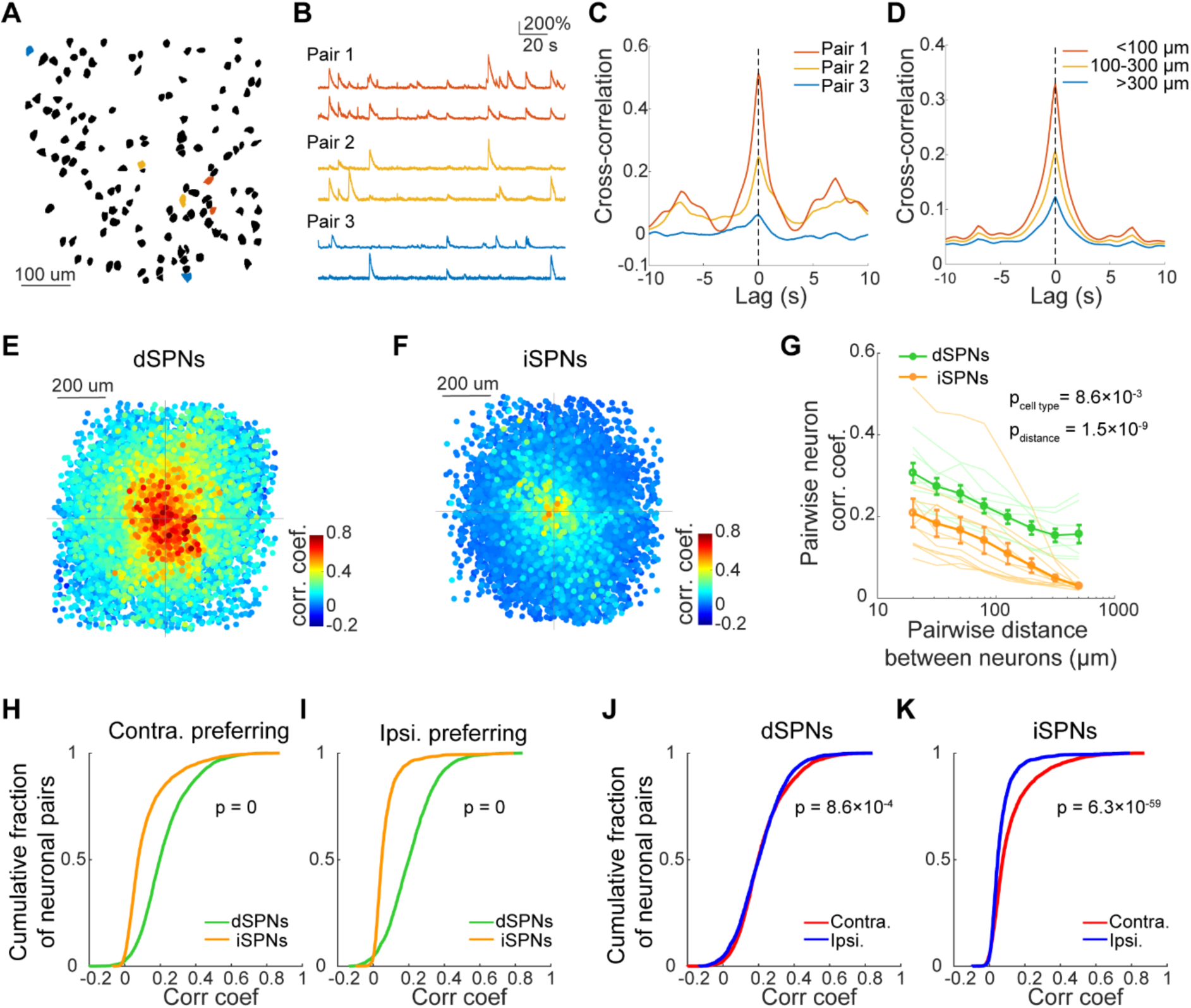
Differential spatiotemporal activity patterns of dSPNs and iSPNs. (A) An example field of dSPNs showing the spatial distribution of regions of interest (ROIs). Colored ROIs indicate pairs of neurons with different spatial distances. Red, <100 μm; yellow, 100-300 μm; blue, >300 μm. (B) Calcium traces from the example neuron pairs shown in (A). (C) Cross-correlation of calcium signals across time for the neuron pairs shown in (B). (D) Cross-correlation across time averaged for all neuron pairs in the example imaging field as in (A) with different distances ranges. Shadings indicate SEM. (E) Two- dimensional view showing spatiotemporal organization of dSPNs from an example imaging field. The coordinate of each dot represents the relative position between a pair of neurons, with one neuron positioned at the origin. The color indicates the correlation coefficients (CCs) between the calcium traces of the corresponding neuron pairs. The third dimension is ordered by the CC values, with higher CC plotted on top layers. (F) Similar as in (E), for an example imaging field of iSPNs. (G) Correlation coefficients between calcium signals of pairs of neurons as a function of the inter-neuronal distance of the corresponding neuron pairs. Each thin line represents data from one imaging field. Green, dSPNs; orange, iSPNs. Data were binned on the dimension of inter-neuronal distance logarithmically scaled for better visualization. Thick lines with error bars, data averaged across imaging fields for dSPNs and iSPNs, respectively. The CCs significantly depend on both the inter-neuronal distance and the cell types, with CCs in dSPNs significantly higher than CCs in iSPNs (Two- way repeated-measures ANOVA, Fisher’s LSD post hoc tests, p _cell type_ = 8.6×10^-3^, p _distance_ = 1.5×10^-9^). (H) Cumulative distribution functions (CDF) of CCs from contralateral-preferring dSPNs and iSPNs (p = 0, two-sample Kolmogorov-Smirnov test, one-tailed). (I) CDF of CCs from ipsilateral-preferring dSPNs and iSPNs (p = 0, two-sample Kolmogorov-Smirnov test, one-tailed). (J) CDF of CCs from contralateral- and ipsilateral-preferring dSPNs (p = 8.6×10^-4^, two-sample Kolmogorov-Smirnov test, one-tailed). (K) CDF of CCs from contralateral- and ipsilateral-preferring iSPNs (p = 6.3×10^-59^, two-sample Kolmogorov-Smirnov test, one- tailed).

We further compared the temporal synchronization in different functional subpopulations of dSPNs and iSPNs with different choice preferences. For both contralateral preferring and ipsilateral preferring subpopulations, the inter-neuronal temporal correlations were also significantly stronger in dSPNs than in iSPNs (**Fig. 3H**, **I**). Within each cell type, the temporal correlations were significantly higher in the contralateral preferring neurons than in ipsilateral preferring neurons for both dSPNs (**Fig. 3J**) and iSPNs (**Fig. 3K**). Thus, consistent with the higher response amplitude in dSPNs than in iSPNs, particularly in the contralateral preferring subpopulations, as shown in **Fig. 2**, dSPNs also show stronger population synchronization than iSPNs. Since higher activity synchrony is likely to produce stronger impact to downstream neurons, the differential temporal synchronization between dSPNs and iSPNs may further contribute to the ensemble competition during decision-making.

### The contralateral preference in choice coding in dSPNs is preserved in the axonal activity in the downstream region

The stronger contralateral response amplitude and the higher temporal synchrony in dSPN ensembles during contralateral trials would concordantly predict a stronger output at the downstream brain regions in contralateral trials. To confirm this, we examined the activity of dSPN axons in the downstream projection target, the substantia nigra pars reticulata (SNr).

The spatial separation between SNr and TS allows us to optically measure the activity of dSPNs axons in SNr without the influence from somatic signals. We expressed GCaMP6s in dSPNs in TS and recorded the axonal calcium signals in SNr using fiber photometry during task performance (**Fig. 4A**; see Methods). We observed reliable axonal calcium signals showing clear preferences for choices (**Fig. 4B**). Across experiments, the dSPN axons in SNr showed significantly stronger responses in contralateral trials than in ipsilateral trials (**Fig. 4C**). Similar to the ROC-based selectivity index in somatic responses as shown in Supplementary Fig. 3, the axonal activity showed significant selectivity to choice directions consistent in correct and error trials (**Fig. 4D**), indicating selective coding for choice but not for sensory information. Across recording sites, the ROC-based selectivity indices were also stronger for the contralateral choices (9/12, p<0.05). Thus, the axonal activity in the output region of basal ganglia also show greater contralateral choice preference, consistent with the contralateral choice preference and temporal synchrony in the soma of dSPNs.

**Figure 4.**
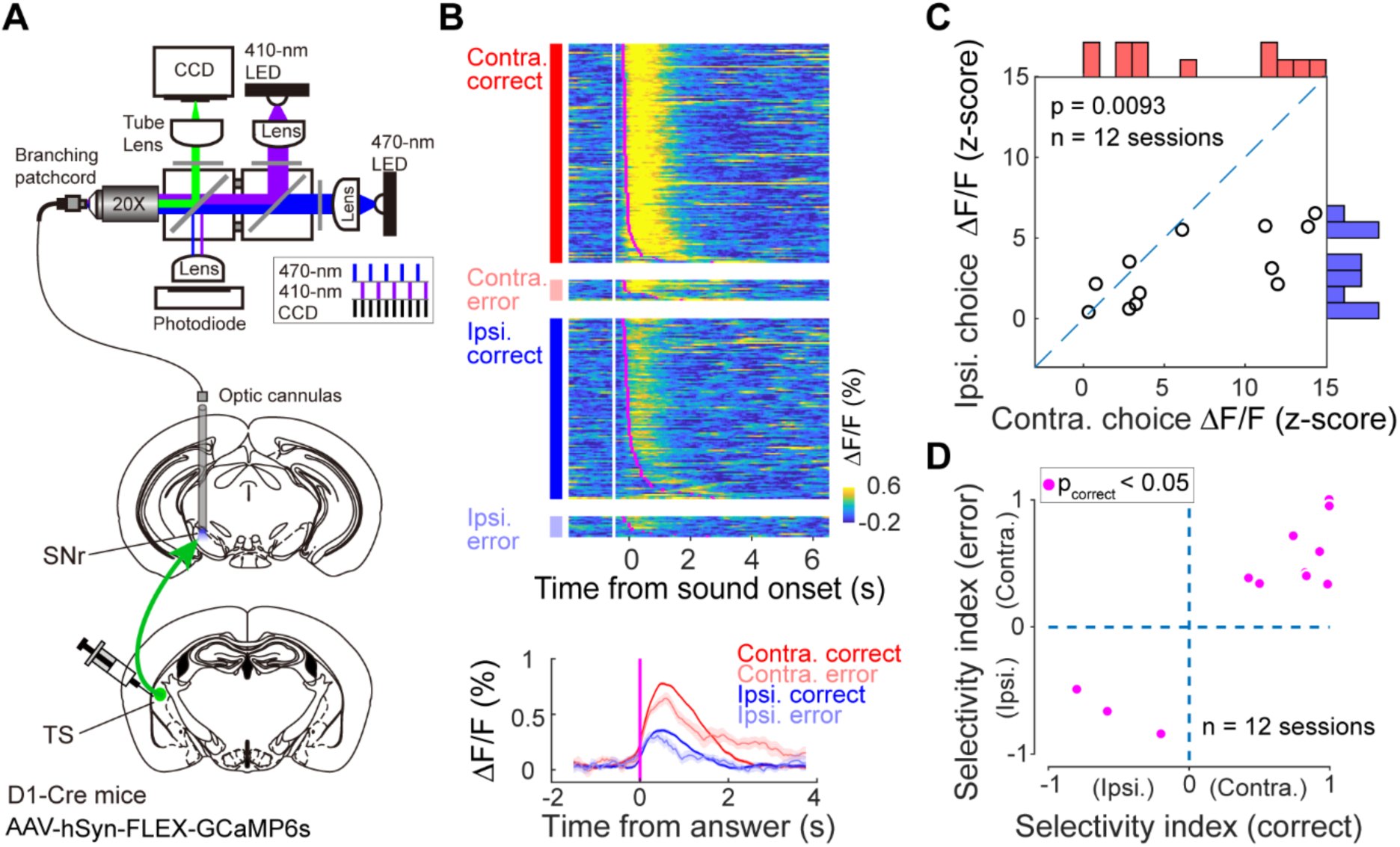
Axonal activity of dSPNs in SNr shows dominant preference for contralateral choices. (A) Experimental configuration of fiber photometry recording of axonal activity of dSPNs in SNr. Top, schematic showing fiber photometry recording system. Bottom, schematic showing virus injection sites in the TS and the fiber implantation in the SNr. (B) Calcium signals from dSPN axons in SNr from an example behavioral session showing greater responses in contralateral trials than in ipsilateral trials. Each row in the color raster plot represents calcium signals from one trial. The calcium signal trials were sorted according to trial types as indicated on the left. White vertical lines, stimulus onset; magenta lines, answer time. Bottom, mean calcium signal traces averaged for the four trial types, aligned to answer times. Shadings indicate SEM. (C) Comparison of axonal activity between contralateral and ipsilateral correct trials. Each dot represents one recording session (n = 12 sessions from 7 mice, p = 0.0093, paired two-sided Wilcoxon signed rank test; see Methods). (D) Selectivity index of dSPN axon activity in SNr for contralateral and ipsilateral trials calculated from correct trials plotted against selectivity index calculated from error trials. Each dot corresponds to one recording session. Magenta dots indicate sessions with significant SI for contra- and ipsilateral choices in correct trials (see Methods).

### Optogenetic activation of the direct and indirect pathways oppositely regulate perceptual decisions

To examine how the information encoded in different subtypes of SPNs impacts perceptual decisions, we examined the causal roles of the striatal SPNs during task performance. First, we performed unilateral optogenetic activation of either dSPNs or iSPNs during the task. To activate dSPNs, we expressed Channelrhodopsin-2 (ChR2) in dSPNs in a Cre-dependent manner by injecting AAV-CAG-FLEX-ChR2 in the TS of D1-Cre mice followed by implanting fiber optics above the injection sites. During experiments, photostimulation (473 nm laser) was delivered to unilateral TS to activate dSPNs on a subset (∼15%) of trials randomly interleaved with control trials without photostimulation (**Fig. 5A** and Supplementary movie 2; see Methods). We found that activation of dSPNs in the TS strongly biased animals’ choices toward the contralateral side relative to the photostimulated hemisphere (**Fig. 5B**, **C**). Our imaging data show that dSPN activity primarily encode choice information rather than sensory information. To confirm that the contralateral bias following dSPN activation was due to an effect on choice rather than on sensory representation, we performed the activation experiments on two separate group of mice trained with reversed sound category to choice direction mappings. One group of mice were required to lick left upon low-frequency sound category and lick right upon high-frequency category. In another group of mice, the sound category to choice mapping was reversed. Activation of dSPNs biased the choice towards the contralateral side in both groups of mice regardless of the associated sound categories (Supplementary Fig. 6), indicating that dSPN activation indeed affected choices rather than sensory representations.

**Figure 5.**
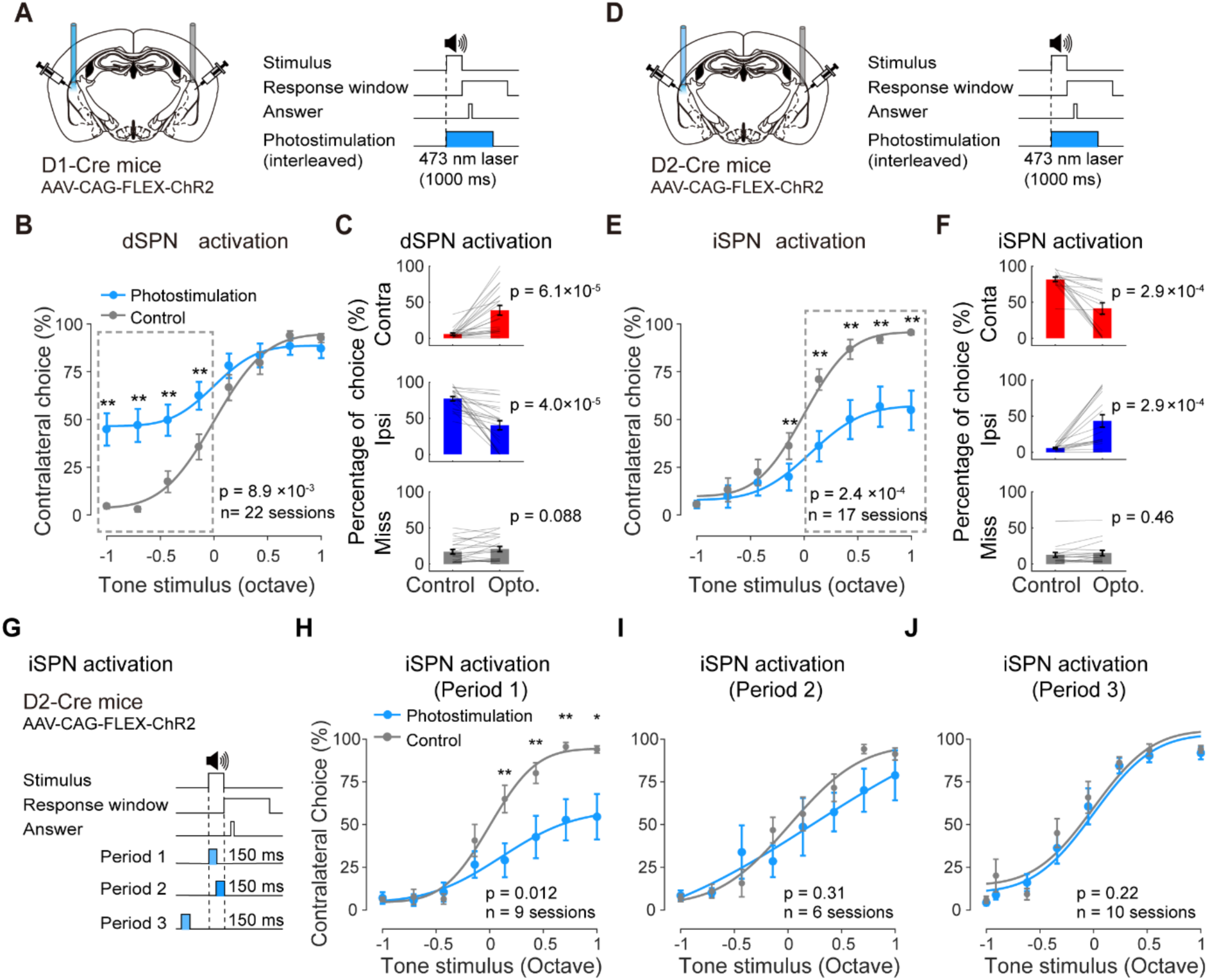
Unilateral optogenetic activation of dSPNs and iSPNs bidirectionally regulate competing choices. (A) Schematic showing experimental configuration for unilateral optogenetic activation of dSPNs. (B) Summarized psychometric functions from control trials (grey) and trials with dSPN activation (blue). Percentage of contralateral choice (relative to photostimulated hemisphere) is plotted against tone stimuli in relative octaves (positive values associated with contralateral choices; negative values associated with ipsilateral choices). n = 22 sessions from 11 D1-Cre mice; p = 8.9×10^-3^ for photostimulation effect, two- way repeated-measures ANOVA; **p < 0.01, Fisher’s LSD post hoc tests. Miss (no lick response) trials were not included. (C) Percentage of choices in trials where tone stimuli are associated with the ipsilateral side as indicated by the negative tone stimulus values in (B). Animals’ responses were compared between control trials and photostimulation trials for contralateral choice (p = 6.1×10^-5^), ipsilateral choice (p = 4.0×10^-5^), and miss (p = 0.088), paired two-sided Wilcoxon signed rank test. (D) Schematic showing unilateral optogenetic activation of iSPNs, similar as in (A). (E) Summarized psychometric functions from control trials and trials with iSPN activation, similar as in (B). n = 17 sessions from 9 D2-Cre mice; p = 2.4×10^-4^ for photostimulation effect, two-way repeated-measures ANOVA; **p < 0.01, Fisher’s LSD post hoc tests. (F) Percentage of choices in trials where tone stimuli are associated with the contralateral side as indicated by the positive tone stimulus values in (E), compared between iSPN activation trials and control trials. Contralateral choice, p = 2.9×10^-4^; ipsilateral choice, p = 2.9×10^-4^; miss, p = 0.46; paired two-sided Wilcoxon signed rank test. (G) Schematic showing optogenetic activation of iSPNs at different periods. Blue shades indicate the period of photostimulation. Period 1, a 150 ms window following sound onset; Period 2, a 150 ms window before the end of tone stimulus; Period 3, a 150 ms window starting from 650 ms before sound onset. (H) Summarized psychometric functions similar as in (E) for iSPN activation in period 1. n = 9 sessions from 6 D2-Cre mice; p = 0.012 for photostimulation effect, two-way repeated-measures ANOVA; *p < 0.05, **p < 0.01, Fisher’s LSD post hoc tests. (I) Similar as in (H) for iSPN activation in period 2. n = 6 sessions from 3 D2-Cre mice; p = 0.31 for photostimulation effect, two-way repeated- measures ANOVA. (J) Similar as in (H) for iSPN activation in period 3. n = 10 sessions from 3 D2-Cre mice; p = 0.22 for photostimulation effect, two-way repeated-measures ANOVA.

Although it was proposed that the dSPN activation could produce positive reinforcement and could change action values [24, 25], the effects on choice behavior here was not due to a persistent change in motivation level since we delivered photonstimulation trial-by-trial in only ∼15% of trials randomly interleaved with control trials precluding potential contributions from accumulated reinforcement. Indeed, the increase of contralateral choices was not accompanied by any significant change in miss rate (**Fig. 5C** and Supplementary Fig. 7B), suggesting that there were no significant changes in motivation level. The effects on choices were neither due to changes in direct motor control of licking, since the lick rate in individual trials was not significantly different between photostimulation and control trials (Supplementary Fig. 9B). Moreover, delivering photostimulation during inter-trial interval did not significantly affect choices (Supplementary Fig. 8). Thus, dSPN activation specifically promotes the decision-related contralateral choices.

To activate iSPNs we injected AAV-CAG-FLEX-ChR2 in the TS of D2-Cre mice (**Fig. 5D**; see Methods). We found that unilateral activation of iSPNs strongly biased animals’ choices toward the ipsilateral side relative to the photostimulated hemisphere (**Fig. 5E**, **F**), an effect opposite to that of dSPN activation. Similar to dSPN activation, we found no significant changes in miss rate during photostimulation (Supplementary Fig. 7C), ruling out potential influence on motivational state by iSPN activation.

Given the general notion that the indirect pathway plays a suppressive role in movement [13, 26], it was unexpected that activation of iSPNs not only reduced contralateral choices but also increased ipsilateral choices without changing the miss rate (**Fig. 5F**). This can be explained by the involvement of the competition between the two pathways and between the two hemispheres during action selection, consistent with the multi-ensemble balancing model inspired by our imaging data (Supplementary Fig. 11).

The precise temporal specificity of photostimulaiton to SPNs during decision-making remains largely unexplored [25,27,28]. We thus examined the temporal specificity of the effects of iSPN activation by applying temporally confined photostimulation during task performance. The time of decision commitment tends to be well before the end of sound stimulus, as we observed that mice often make correct answer lick before the end of sound stimulus (∼150 ms after sound onset, Supplementary Fig. 10A). We thus restricted the photoactivation of iSPNs within a 150 ms window after sound stimulus onset (the first half of sound period), covering the decision period while excluding most answer licks (**Fig. 5G**, period 1). We found that this precisely confined photoactivation also strongly biased animals’ choice toward the ipsilateral side (**Fig. 5H**). Next, we restricted the photoactivation to the second half (150 ms) of the sound period contained most post-decision answer licks (**Fig. 5G**, period 2). The choice behavior was largely unaffected following this period 2 photoactivation (**Fig. 5I**). In addition, when delivering the photoactivation in a 150 ms window during the inter-trial interval before the sound stimulus onset (**Fig. 5G**, period 3), we found no significant changes in the behavioral performance either (**Fig. 5J**). Together, these results suggest that rather than having a general suppressive effect on movement, iSPNs can be precisely involved the decision process before motor execution.

### Optogenetic inactivation reveals opposite contributions from dSPNs and iSPNs to choice behavior

Previous studies on the causal roles of dSPNs and iSPNs during choice behavior mainly focused on optogenetic activation [25, 27]. To understand more physiological contributions of dSPNs and iSPNs to decision-making, we performed optogenetic inactivation of dSPNs and iSPNs during task performance. To inactivate dSPNs, we injected AAV-Ef1α-DIO- eNpHR3.0 in the TS of D1-Cre mice, and delivered unilateral photostimulation (593 nm) through fiber optics on ∼15% interleaved trials (**Fig. 6A**). We found that inactivation of dSPNs significantly biased animals’ choice toward the ipsilateral side relative to the photostimulated hemisphere comparing to control trials (**Fig. 6B**, **C**, *P* = 3.8×10^-3^, two-way repeated-measures ANOVA). To inactivate iSPNs, we injected AAV-Ef1α-DIO-eNpHR3.0 in the TS of D2-cre mice (**Fig. 6D**). We found that iSPN inactivation led to a slight but significant bias of animals’ choice toward the contralateral side (**Fig. 6E** and **6F**, *P* = 1.7×10^-4^, two-way repeated-measures ANOVA), an opposite effect comparing to that by dSPN inactivation. To confirm the relatively weak but significant behavioral effects, especially following iSPN inactivation, we expressed the soma-targeted *Guillardia theta* anion-conducting channelrhodopsin-2 (stGtACR2), a more potent activity suppressor [29], in iSPNs, and delivered blue laser (473 nm) to inactivate iSPNs during task performance (**Fig. 6G**). We found significant contralateral bias following iSPN inactivation by stGtACR2 (**Fig. 6H**, **I**), similar to that following inactivation using eNpHR3.0. Unilateral inactivation of dSPNs or iSPNs did not influence the miss rate or lick rate (Supplementary Fig. 7D-I, Supplementary Fig. 9D-F), ruling out potential effects on motivation or tongue movement.

**Figure 6.**
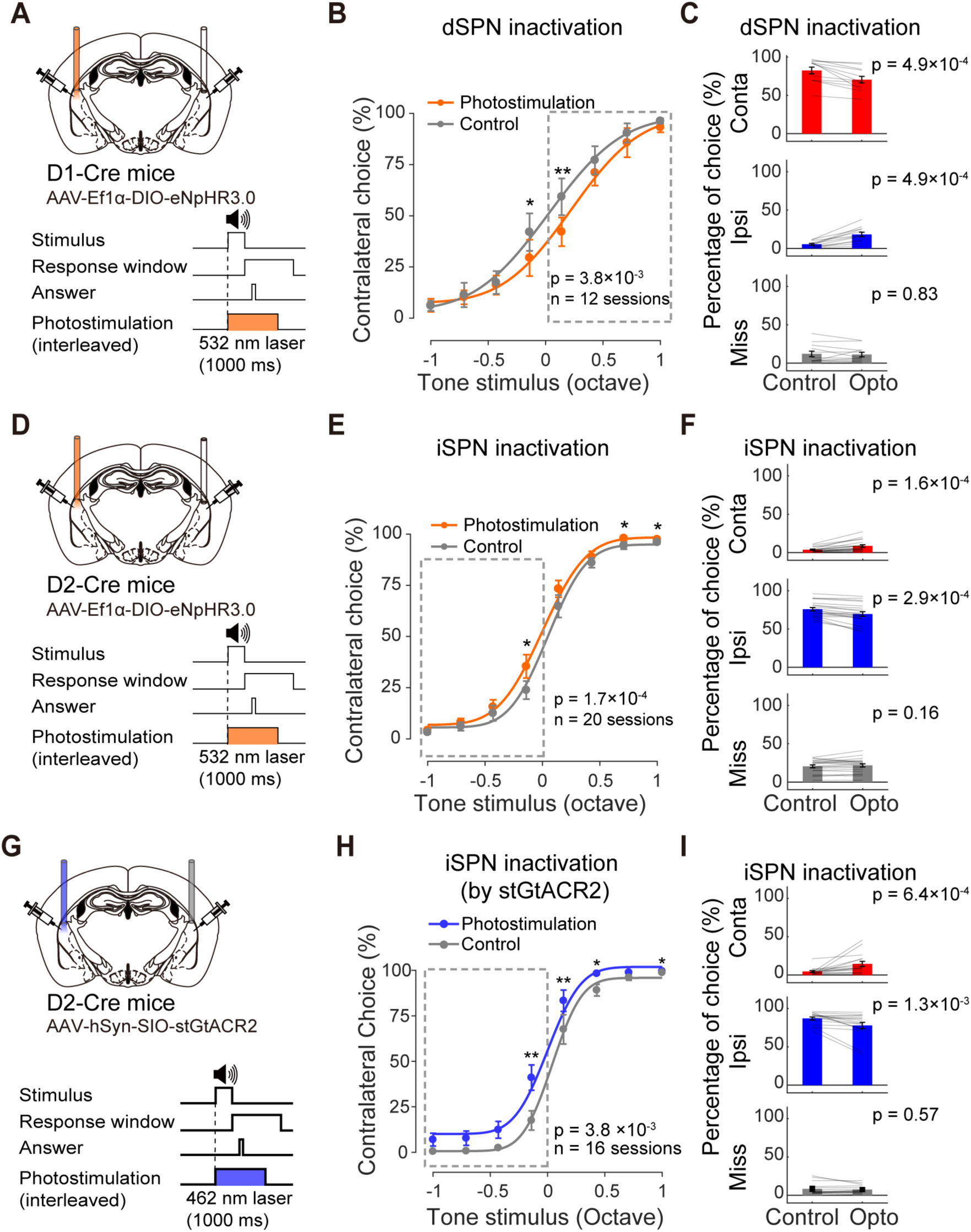
Unilateral optogenetic inactivation of dSPNs and iSPNs supports their complementary contributions to behavioral choices. (A) Schematic showing experimental configuration for unilateral optogenetic inactivation of dSPNs using eNpHR3.0. (B) Summarized psychometric functions for unilateral dSPN inactivation. n = 12 sessions from 6 D1-Cre mice; p = 3.8×10^-3^ for photostimulation effect, two-way repeated-measures ANOVA; *p < 0.05; **p < 0.01, Fisher’s LSD post hoc tests. (C) Percentage of choices in trials where tone stimuli are associated with the contralateral side as indicated by the positive tone stimulus values in (B). Animals’ responses were compared between control trials and photostimulation trials for contralateral choice (p = 4.9×10^-4^), ipsilateral choice (p = 4.9×10^-4^), and miss (p = 0.83), paired two-sided Wilcoxon signed rank test. (D) Schematic for unilateral iSPN inactivation using eNpHR3.0. (E) Summarized psychometric functions for unilateral iSPN inactivation. n = 20 sessions from 10 D2-Cre mice; p = 1.7×10^-4^ for photostimulation effect, two-way repeated-measures ANOVA; *p < 0.05, Fisher’s LSD post hoc tests. (F) Percentage of choices in trials where tone stimuli are associated with the ipsilateral side as indicated by the negative tone stimulus values in (E), compared between iSPN inactivation trials and control trials. Contralateral choice, p = 1.6×10^-4^; ipsilateral choice, p = 2.9×10^-4^; miss, p = 0.16; paired two-sided Wilcoxon signed rank test. (G) Schematic for unilateral iSPN inactivation using stGtACR2. (H) Summarized psychometric functions for unilateral iSPN inactivation using stGtACR2. n = 16 sessions from 8 D2-Cre mice; p = 3.8×10^-3^ for photostimulation effect, two-way repeated-measures ANOVA; *p < 0.05, **p < 0.01, Fisher’s LSD post hoc tests. (I) Percentage of choices in trials where tone stimuli are associated with the ipsilateral side as indicated by the negative tone stimulus values in (H), compared between iSPN inactivation trials (by stGtACR2) and control trials. Contralateral choice, p = 6.4×10^-4^; ipsilateral choice, p = 1.3×10^-3^; miss, p = 0.57; paired two- sided Wilcoxon signed rank test.

Thus, unilateral inactivation of dSPNs or iSPNs revealed opposing contributions to choice behavior, consistent with the expectations based on our imaging and activation results.

Together, our imaging and manipulation results show that multiple ensembles of dSPNs and iSPNs exhibit differential response strengths and play opposing causal roles in selecting either contralateral or ipsilateral actions during decision behavior. These ensembles from both hemispheres therefore form a multi-component balancing system that could be tightly regulated and flexibly adjusted to actuate decision-related action selection (Supplementary Fig. 11).

### Concurrent disinhibition of dSPNs and iSPNs promotes contralateral choice

To test the whether the differential response patterns between dSPNs and iSPNs would contribute to action selection as suggested in our multi-ensemble balancing model (Supplementary Fig. 11), we sought to concurrently activate both pathways while preserving their endogenous activity patterns. Since both dSPNs and iSPNs receive comparable inhibitory input from local parvalbumin (PV) expressing interneurons [30–33], we reasoned that inhibiting the PV interneurons would mimic coactivation of both dSPNs and iSPNs with intact endogenous excitatory input patterns. We first examined whether striatal PV interneurons show any task-related activity by recording their calcium signals using fiber photometry during task performance. We expressed GCaMP6s in PV interneurons by injecting AAV-hSyn-FLEX-GCaMP6s in the TS of PV-Cre mice, and implanted fiber optics above the injection sites (**Fig. 7A**). We found that PV interneurons showed strong task modulated responses with clear alignment to choices (**Fig. 7B**). These responses are comparable in contra- and ipsilateral trials (**Fig. 7C**), suggesting that their overall excitatory inputs representing opposing choices are balanced. Therefore, experimentally inhibiting PV interneurons would disinhibit both subtypes of SPNs without introducing additional biasing factors.

**Figure 7.**
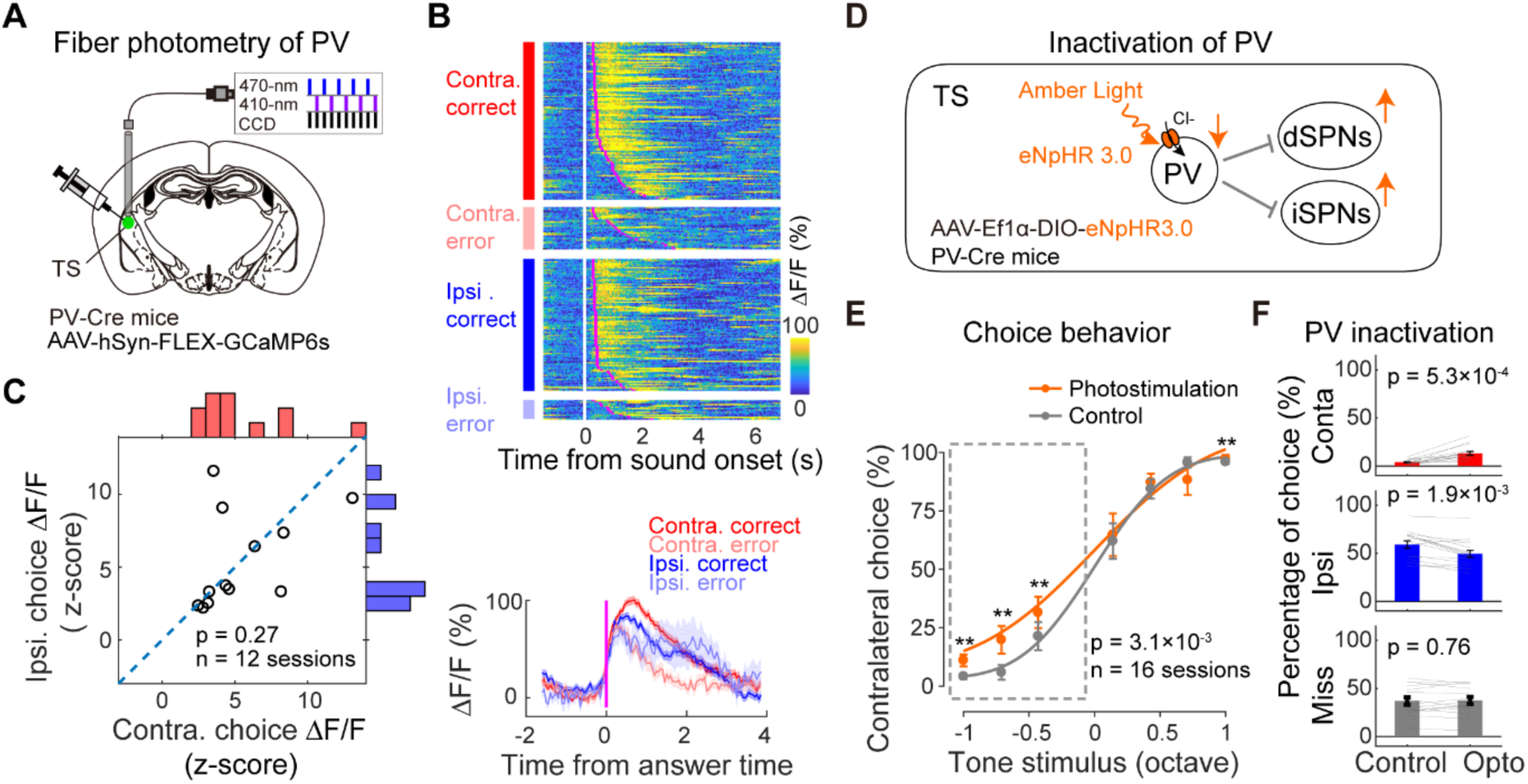
Concurrent disinhibition of dSPNs and iSPNs biases choices towards the contralateral side. (A) Schematic showing fiber photometry recording from PV interneurons in TS. (B) Calcium signals of PV interneurons in the TS from an example behavior session. Upper, color raster plot of calcium signals from individual trials sorted according to the four trial types; lower, mean traces averaged for different trial types (Similar as in Fig. 4B). (C) Summarized responses of PV interneurons compared between trials with contralateral and ipsilateral choices. n = 12 sessions from 6 mice, p = 0.27, paired two-sided Wilcoxon signed rank test. (D) Schematic showing optogenetic inactivation of PV interneuron in TS. (E) Summarized psychometric functions following unilateral PV interneuron inactivation. n = 16 sessions from 8 PV-Cre mice; p = 3.1×10^-3^ for photostimulation effect, two-way repeated-measures ANOVA; **p < 0.01, Fisher’s LSD post hoc tests. (F) Percentage of choices in trials where tone stimuli are associated with the ipsilateral side as indicated by the negative tone stimulus values in (E), compared between PV inactivation trials and control trials. Contralateral choice, p = 5.3×10^-4^; ipsilateral choice, p = 1.9×10^-3^; miss, p = 0.76; paired two- sided Wilcoxon signed rank test.

We thus performed optogenetic inactivation of the striatal PV interneurons by expressing eNpHR3.0 in these neurons using AAV-Ef1a-DIO-eNpHR3.0 in PV-Cre mice (**Fig. 7D**). We found that unilateral inactivation of PV interneurons in the TS significantly biased choices toward the contralateral side (**Fig. 7E**, **F**). This result is consistent with the prediction from the multi-ensemble balancing model that coordinated activation of both dSPNs and iSPNs support contralateral action selection following decision-making (Supplementary Fig. 11).

### Computational implementation of the coordination among SPN ensembles

Based on our imaging data (**Fig. 2**), both dSPNs and iSPNs comprise divergent functional subpopulations with response preferences to competing choices. The competition and cooperation among these functional subpopulations from both hemispheres would determine the final behavioral choices. To explore the computational plausibility of such coordination mechanism, we constructed a network model of the basal ganglia circuitry incorporating the multiple functional subpopulations of SPNs in the two hemispheres (**Fig. 8A**). In this model, the sensory decisions are derived from cortex and sent to dSPNs, iSPNs and striatal PV neurons via cortico-striatal projections [34]. Within the local striatal region, PV neurons provide inhibitory input to both dSPNs and iSPNs, and the two subtypes of SPNs provide mutual inhibitory input to each other [30,31,33,35]. The dSPN and iSPN populations provide either inhibitory or excitatory (indirectly) input to ipsilateral SNr through the direct and indirect pathways respectively [11, 36]. The left and right SNr populations compete with each other, and the dominant SNr population determines the left or right choices. The weights of cortical input to different subpopulations of SPNs were set according to their relative response strengths observed in our imaging experiments (Supplementary table 1). With these settings, we recapitulated the relative response strengths found in our experimental data using our model (**Fig. 8B**, **C**).

**Figure 8.**
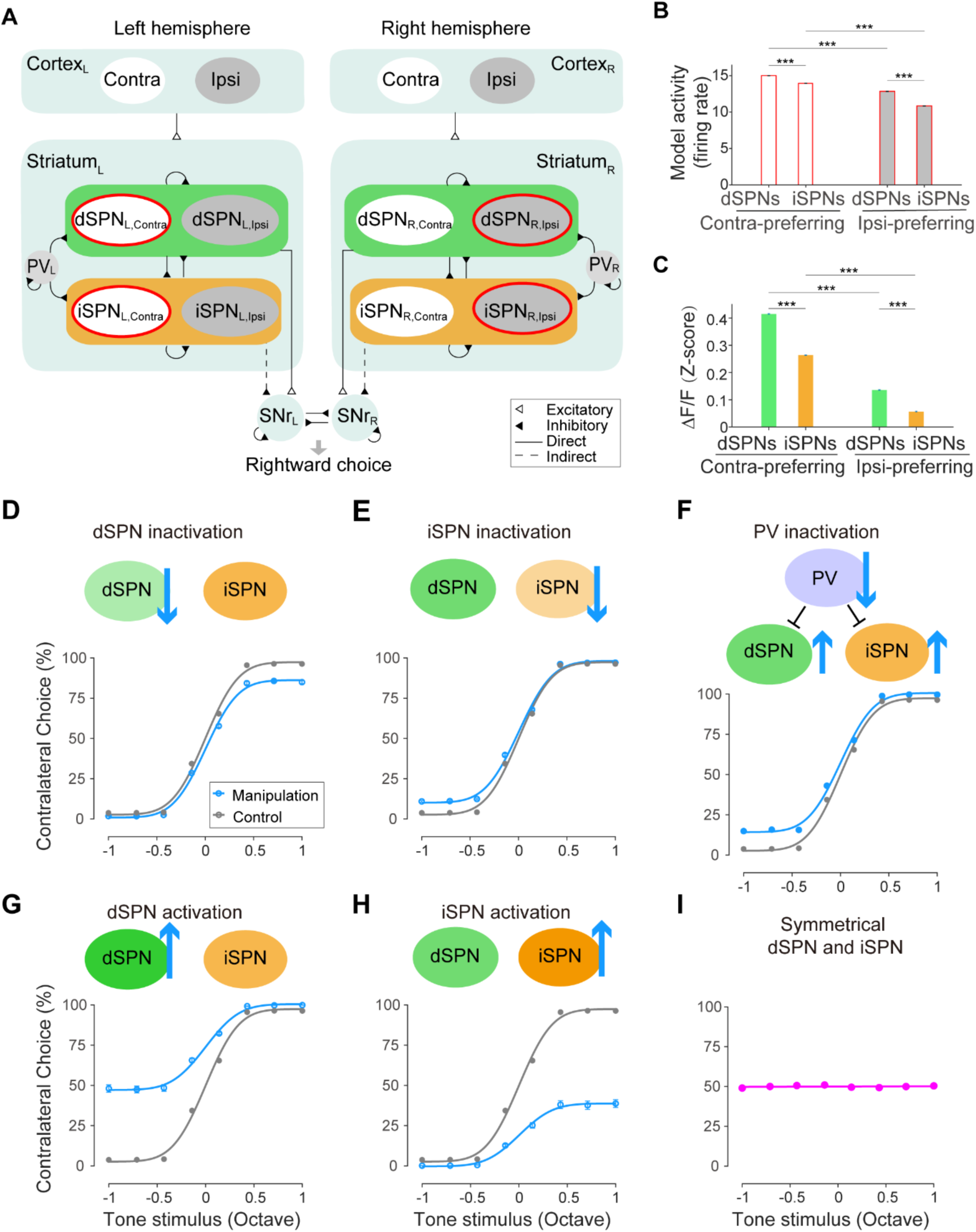
A neural network model recapitulates the causal roles of the dSPNs and iSPNs in perceptual decision behavior. (A) Schematic showing the network model organization. L, left hemisphere; R, right hemisphere; Contra, contralateral preferring neuronal population; Ipsi, ipsilateral preferring neuronal population. See Methods. (B) Responses of modeled dSPNs and iSPNs with contralateral or ipsilateral preference in preferred trial types. ***, p <0.001, two-sided Wilcoxon rank sum test. (C) Experimental data of dSPN and iSPN responses same as in Fig. 2Q. (D-H) Model simulation of behavioral effects of optogenetic manipulations. Model outputs of choice behavior from control trials were shown in grey, and those from trials with unilateral dSPN activation (D), iSPN activation (E), dSPN inactivation (F), iSPN inactivation (G) and PV interneuron inactivation (H) were shown in blue. Each curve was obtained from 22 different *I ^stim^* values (Supplementary table 2). (I) Simulated choice behavior when dSPN and iSPN activity are symmetrical.

To examine whether the coordination of multiple SPN ensembles could be computationally implemented to produce expected choice behavior, we asked whether our network model could recapitulate the perceptual decision behavior under both control condition and cell-type specific manipulation conditions. First, simulation using this model was able to recapitulate the choice behavior with similar psychometric functions as in behavioral data (**Fig. 8D**-**H**). Importantly, when we removed the difference between dSPNs and iSPNs *in silico*, the model was no longer able to produce the choice behavior based on sensory stimuli (**Fig. 8I**), corroborating the importance of the difference between dSPN and iSPN ensembles found in our imaging results in supporting choice behavior. Next, we used this model to simulate optogenetic manipulations by increasing or decreasing the activity level of different striatal cell types (Supplementary table 2). We found that unilateral increasing of dSPN activity biased the choice towards the contralateral side, whereas unilateral increasing of iSPN activity biased the choice towards the ipsilateral side (**Fig. 8D**, **E**), recapitulating our experimental results of optogenetic activation of dSPNs and iSPNs (**Fig. 5A**-**F**). Conversely, unilateral decreasing dSPN activity significantly biased the choice towards ipsilateral side, whereas unilateral decreasing iSPN activity significantly biased the choice towards contralateral side (**Fig. 8G**-**H**), which also recapitulated our optogenetic inactivation results (**Fig. 6**). Finally, we simulated the concurrent disinhibition by decreasing the activity of unilateral PV interneurons, which provide inhibitory input to both dSPNs and iSPNs with similar connectivity weights in the model (Supplementary table 1 and table 2). Decreasing unilateral PV interneuron activity biased the choice towards the contralateral side (**Fig. 8F**), recapitulating our experimental observation following PV neuron inactivation (**Fig. 7E**). These simulation results demonstrate the plausible computational implementation of the coordination among SPN ensembles with the differential response patterns between dSPNs and iSPNs as shown in our experimental data, which supports the perceptual decision-making behavior.

## Discussion

It has been a long-standing debate how the two opposing basal ganglia pathways compete and cooperate to elicit desired actions. This problem is particularly critical for decision-making where proper actions need to be selected from competing motor plans. By combining cell-type specific *in vivo* two-photon imaging and optogenetic manipulations with a psychophysical decision-making task, we show that the posterior tail of the striatum contains functionally divergent subpopulations of SPNs forming a multi-ensemble balancing system (Supplementary Fig. 11), which plays an essential role in sensory-guided action selection. Our single-cell resolution imaging reveals that both dSPNs and iSPNs contain subpopulations with divergent preferences to competing choices, consequentially resulting in concurrent activation of different ensembles of both dSPN and iSPN in response to the same motor choices (**Fig. 2**). This is consistent with previously observed coactivation of the direct and indirect pathways during movement. Importantly, our imaging data also reveal that the dSPNs and iSPNs show ensemble-level difference in their contralateral choice preference during decision-making, which was essential in driving the decision execution. Corroborating the differences in response amplitude, dSPNs show higher level of population synchronization than iSPNs (**Fig. 3**), likely producing stronger impact on downstream neurons. These imaging results inspire a multi-ensemble coordination mechanism for selecting competing actions during decision-making (Supplementary Fig. 11).

Cell-type specific optogenetic manipulations of dSPNs and iSPNs reveal opposing effects on choice behavior with the temporal specificity in the decision-relevant period (**Fig. 5** and **Fig. 6**). These manipulation results extend the classical notion of the differential roles of the direct and indirect striatal pathways in movement control to the domain of perceptual decision-making with trial-by-trial specificity and sub-trial precision. Moreover, concurrent optogenetic disinhibition of dSPNs and iSPNs led to significant contralateral bias of choice behavior (**Fig. 7**), suggesting that physiological level coactivation of dSPNs and iSPNs can drive desired motor choices, supporting the multi-ensemble coordination model (Supplementary Fig. 11). Finally, a neural circuit model featuring the multi-ensemble structure recapitulated both the perceptual decision-making behavior and the behavioral effects following cell-type specific optogenetic manipulations (**Fig. 8**), supporting plausible computational implementation of our multi-ensemble coordination model.

Although SPNs in the TS receive synaptic input from auditory regions, we found that only a small number of neurons showed sensory selectivity, while the majority of neurons of both dSPNs and iSPNs showed strong selectivity for choices (Supplementary Fig. 3). This can be partially due to that the calcium signals detected using GCaMP6 primarily reflect strong spiking activity related to decision readout. We verified this coding preference for choice information by recording the axonal activity of dSPNs using fiber photometry in the downstream SNr and also found primarily choice selective responses with stronger contralateral preference (**Fig. 4**). This is also consistent with a previous study using fiber photometry to record GCaMP6 signals from dSPNs during a visual detection task showing predominantly licking-related signals instead of visual responses [27].

Recent studies reported coactivation of both the dSPNs and iSPNs during movement [16–20]. These observations appear to contradict predictions from earlier models derived from the opposing effects on movement of the direct and indirect pathways [4,10,13]. Such discrepancy could be potentially reconciled if there were task-specific differentiation in the activity patterns between dSPNs and iSPNs [4,15,37,38]. Here we found that dSPNs and iSPNs indeed exhibit ensemble-level differences in the contralateral dominance during choice behavior, with stronger contralateral population responses in dSPNs beginning prior to decision execution (**Fig. 2**). Meanwhile, the divergent subpopulations in both dSPNs and iSPNs found in our data implies that for a given choice, subpopulations from both pathways could be co-activated. Such ensemble organization would explain the concurrent activation of dSPNs and iSPNs, while also support selecting the proper actions during decision-making (Supplementary Fig. 11).

In agreement with this possibility, a previous study showed that concurrent over- activation of both direct and indirect striatal pathways biased the animals’ locomotion towards the contraversive direction [20]. Here, instead of over-activation, we tested whether co-activation of both pathways with endogenous excitatory inputs would promote contralateral choices during decision-making. To this end, we concurrently disinhibited both pathways by optogenetic inactivation of PV interneurons, a common source of inhibition to dSPNs and iSPNs. This would effectively co-activate the two pathways by their endogenous excitatory inputs. Indeed, this manipulation significantly biased animals’ choice towards the contralateral side (**Fig. 7**), consistent with the multi-ensemble coordination model inspired by our imaging data. In accordance, concurrently activating dSPNs and iSPNs in our neural circuit model also biased the choice toward the contralateral side (**Fig. 8**). Notably, this simulation result depends on the prerequisite of the differential contralateral preferences between dSPNs and iSPNs. Thus, our results demonstrate that the concurrent activation of dSPNs and iSPNs with differential response preferences can effectively select desired actions during decision-making.

In the past, various models have been proposed to account for the role of the striatal pathways in movement control [4,11,13,15,37,39]. Our results extend the classical rate model, which primarily concerns movement initiation, to decision-making related action selection, by providing a multi-ensemble coordination mechanism in the direct and indirect pathways. In this sense, our results are also largely consistent the “competitive” model [37].

Other models, such as the “complementary” model [11,28,37,39] concerning a role of the indirect pathway in suppressing unselected movements outside the decision-making task, may be further tested in future studies examining both task and non-task related motor actions.

The causal roles of the cell-type specific striatal circuits have been widely studied in various forms of motor function, including locomotion [14, 19], sequential actions [28, 40], and movement kinematics [41]. For sensory-guided decision-making, however, the causal roles of specific striatal pathways are less understood, especially regarding their relative contributions to decision process versus ensuing movements. We found that activation of dSPNs and iSPNs elicited strong but oppositely lateralized decision-bias (**Fig. 5**), consistent with the general notion of the functional opponency of the direct and indirect striatal pathways. By precisely restricting photostimulation in different 150 ms windows, we demonstrate that iSPN activation contribute to action selection specifically during decision process before motor execution. To our knowledge, this is the first demonstration distinguishing the causal role of striatal circuits in decision process from movement control. In addition to activation, it is important to also examine the effect of inactivation to reveal the causal involvement of neuronal activity. Previous studies mainly focused on optogenetic activation [25, 27]. Here, we found that optogenetic inactivation of dSPNs and iSPNs led to opposite lateralized biasing of choice behavior with reversed directions than the activation effects (**Fig. 6**), revealing the causal involvement of the two striatal pathways in decision-making behavior.

A recent study on a visual detection go/no go task showed that, intriguingly, activation of dSPNs and iSPNs did not lead to opposite actions, but both increased licking upon visual stimulation, although the effect was stronger for dSPN activation when the visual stimulus was on the contralateral side [27]. This might be partially attributable to the behavioral readout of the go/no-go task not distinguishing lateralized actions. This issue is largely resolved in our two-choice decision-making task, allowing for disambiguating the distinct lateralized causal roles for both dSPNs and iSPNs.

The differential ensemble response patterns between dSPNs and iSPNs could be attributable to the differential synaptic strength in the cortical or thalamic input to different subtypes of SPNs, or could also be due to lateralized dopaminergic input [42], and may result from dopamine-dependent plasticity [43]. The mechanism for the regulation of synaptic input from the cortical and thalamic neurons encoding different sensory and motor variables during behavior and learning is an important subject to be further investigated in the future.

## Methods

Information on materials used to conduct the research, and all methods used in the analysis are available in the supplementary data.

## Acknowledgements

We thank Dr. Mu-ming Poo, Dr. Yong Gu, Dr. Yangang Sun and Dr. Chengyu Li for valuable discussions; Dr. Chunyu Ann Duan and Dr. Yu Xin for help in data analysis; Mengjun Sheng and Di Lu for help in surgery; Zhaomei Ying for lab assistant.

## Funding

This work was supported by the National Key R&D Program of China (No. 2021YFA1101804); National Science and Technology Innovation 2030 Major Program (No. 2021ZD0203700 / 2021ZD0203704); Shanghai Pilot Program for Basic Research – Chinese Academy of Science, Shanghai Branch (No. JCYJ-SHFY-2022-011); Lin-Gang Lab (Grant No. LG2021XX-XX-XX LG202104-01-05); Gift Funding Project on Prior Knowledge-based Artificial Neural Network Research, Huawei RAMS Technologies Lab; the National Science Fund for Distinguished Young Scholars (to N.L.X.).

## Author contributions

L.C., S.T. and N.L.X. conceived the project and designed the experiments. L.C. performed the optogenetic experiments. S.T. developed the striatum chronic imaging window method and performed the two-photon imaging experiments. J.P. designed the fiber photometry system. J.P. and N.L.X. developed the behavioral system. K.Z. and B.S. constructed the neural network model and performed the model simulation. Z.Z. performed part of the behavioral training and optogenetic experiments. L.C., S.T. and N.L.X. analyzed the data and wrote the manuscript.

## Declaration of interests

The authors declare no competing financial interests.

## Materials & Correspondence

Correspondence and requests for materials should be addressed to N.L.X.

## Data and materials availability

All data to understand and assess the conclusions of this study are available in the main text or supplementary materials. All the original behavioral, imaging, histochemical data and analysis code are archived in the Institute of Neuroscience, CAS Center for Excellence in Brain Science and Intelligence Technology, Chinese Academy of Sciences.

## Methods

### Subjects

All experimental aspects were carried out in compliance with the procedures approved by the Animal Care and Use Committee of the Institute of Neuroscience, Chinese Academy of Sciences (Center for Excellence in Brain Science and Intelligence Technology, Chinese Academy of Sciences). Male mice, 2- to 3-month-old, D1-Cre (MMRRC_030989-UCD) and D2- Cre (MMRRC_032108-UCD) backcrossed into Black C57BL/6J were used. Male mice, 2- to 3- month-old Black C57BL/6J and PV-Cre mice (Jax stock #008069) were also used. Mice had no previous history of any other experiments. Mice used for behavioral experiments were housed in a room with a reverse light/dark cycle. On days without behavioral training or testing, the water-restricted mice received 1 mL of water. On experimental days, mice receive 0.5 to 1 mL of water during each behavioral session which lasted for 1 to 2 hours. Body weight was measured daily, and was maintained above 80% of the weight before water restriction.

### Virus injection for two-photon imaging

D1-Cre or D2-Cre mice were anesthetized with isoflurane (1∼2%) mixed in air (Pin-Indexed fill or Funnel-Fill Isoflurane Vaporizer, VETEQUIP), and placed on a stereotaxic frame (RWD 68043 and 68077). Throughout the surgery, body temperature was monitored and maintained at 37 ℃ by temperature controller (RWD 69002), and eyes were moisturized with eye ointment. After shaving the hair, the skin was sterilized with 75% ethanol and iodine. A piece of skin was removed to expose the skull. After cleaning the skull with a scalpel removing connective tissues, a small craniotomy (around 0.5 mm in diameter) was made on the skull above the TS (AP: -1.46, ML: 3.25) for virus injection. The coordinate for cannular window implantation was also marked (AP: -1.0, ML: 4.7). Cre-dependent GCaMP6s virus (200-300 nL AAV2/9-hSyn-FLEX-GCaMP6s) was injected slowly with a hydraulic manipulator (Narashige, MO-10) through a glass micropipette (pulled using Sutter Instrument P-97, with tip broken into an opening with 20-30 μm inner diameter; Drummond Scientific, Wiretrol II Capillary Microdispenser) into the TS at a depth of -3.4 mm from the level of bregma.

After injection, the micropipette was kept in the injection site for ∼10 min before being retracted slowly to reduce flow back of the viruses. The craniotomy was then covered with silicon adhesive (Kwik-Sil, WPI). The exposed skull and surrounding tissues was then covered with instant adhesive (LOCTITE 495, Henkel). A custom-designed titanium headplate was then attached to the skull with dental cement for head fixation during behavioral sessions. After surgery, mice were allowed to recover for at least 7 days before water restriction and behavioral training. Virus preparations were obtained from Shanghai Taitool Bioscience Co.Ltd, or OBiO Technology (Shanghai) Corp., Ltd.

### Behavioral training

After one-week recovery following surgery, mice were started with water restriction (1 mL per day), which lasted ∼7 days before the beginning of behavior training. Mice performed one behavioral training session each day, and were allowed to perform the task until sated. Body weights were measured and water consumption was estimated using the body weight difference before and after each session. If mice consumed less than 0.5 mL water per day, additional water supplement was provided.

The behavioral apparatus has been described previously [1, 2]. Briefly, experiments were conducted inside custom-designed double-walled sound-attenuating boxes. The mouse auditory decision behavior was controlled by a custom-developed PX-Behavior System. Cosine ramps (5 ms) were applied to the rising and falling of all tone stimuli. The sound waveforms are amplified using ZB1PS (Tucker-Davis Technologies) and delivered by an electrostatic speaker (ES1, Tucker-Davis Technologies) placed on the right side of the animal. The sound system was calibrated using a free-field microphone (Type 4939, Brüel and Kjær) over 3–60 kHz to ensure a flat spectrum (±5 dB SPL). Behavioral trials were synchronized with two-photon image acquisition by digital outputs from the PX-Behavior System to the ‘ScanImage’ software which controls the two-photon imaging setup. In optogenetics experiments, the lasers were also controlled through digital outputs from PX-Behavior System.

The behavioral task is based on the auditory-guided two-alternative-forced-choice (2AFC) task on head-fixed mice as described before [1, 2]. After mice learnt how to lick left and right water ports, they were trained to discriminate a range of tone frequencies (between 7 and 28 kHz, or between 5 and 20 kHz, distributed in logarithmic scale of octaves) as higher or lower than the defined category boundary, which is the mid line of the logarithmically spaced frequency range, i.e., 14 kHz for the 7-28 kHz range, and 10 kHz for the 5-20 kHz. The frequency ranges were chosen according to the typical hearing range of laboratory mouse [3]. Sound intensity was 70 dB SPL for all used frequencies. The initiation of each trial is not explicitly cued, and the animal needs to wait for the tone stimulus to occur. Following the inter-trial interval (ITI), a 0.5-1 s random delay was imposed before tone stimulus (duration, 300 ms) to ensure that animals cannot predict the onset of tone stimuli. Mice were required to respond by licking left or right water port within a 3 s answer period following the stimulus offset to report their perceptual judgements of the tone as the high or low frequency categories. Correct responses led to opening of the water valve for a short period of time to dispense a small amount of water reward (∼1.5 μL). Error responses lead to a 2∼6 s time-out punishment, during which licking to the wrong side would reinitiate the time-out period. To control for systematic side bias, different groups of mice were trained with opposite stimulus-response contingencies, i.e., high-left / low-right, or high-right / low-left (Supplementary Fig. 1 B-C and Supplementary Fig. 6A). Mice completed several hundred trials in each behavioral session allowing for sufficient statistical power quantifying perceptual decisions on stimulus categorization. Mice were first trained with the two easy frequencies separated by 2 octaves, e.g., 7 and 28 kHz, or 5 and 20 kHz until reach 85% correct rate (after ∼10 session, Supplementary Fig. 12), after which 6 tone stimuli of intermediate frequencies (between 7 and 28 kHz or 5 and 20 kHz, linearly spaced on the logarithmic scale) were delivered in randomly interleaved trials. For each session, the fraction of trials with tones of the intermediate frequencies is ∼30% to keep the performance stable. The trial numbers in each imaging or optogenetic session were showed in Supplementary Fig. 13.

Psychometric function was obtained by fitting the behavioral data using a 4-parameter sigmoidal function [4, 5]

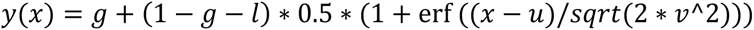

where *y*(*x*) is the probability that animal would make a right choice (**Fig. 1C**) or a contralateral choice, and *x* is the tone frequency (in octave). Parameters to be fitted are *g* the guess rate, *l* the lapse rate, *u* the subject bias (boundary), and *v* the discrimination sensitivity (threshold). erf () represents error function. For psychometric function fitting, ‘miss’ trials in which mice did not response during the answer period were excluded. To quantify the proportion of contralateral choices in optogenetic manipulation and imaging experiments, we pooled all animals trained with either of the two types of contingencies (‘low-left / high-right’ or ‘low-right / high-left’) by ordering the tone frequencies according to relative octaves distance from the boundary frequency, with negative values representing frequencies associated with ipsilateral choices and positive values the contralateral choices.

### Cannula window implantation for two-photon calcium imaging

When the behavioral performance reached the criterion of 85% correct, mice were allowed to have free access to water for 2-3 days before the surgery of implanting cannular imaging window. During the surgery, mice were anaesthetized with isoflurane (1∼2%), and an ∼2 mm diameter craniotomy was made centered at the marker for window implantation. The tissue above the TS were removed by aspiration through 25-gauge and 27-gauge blunt needles connected to a peristaltic pump (LongerPump YZ1515x). During the aspiration, saline was continuously delivered, and moisturized gelatin sponge (Xiang’en) was applied to prevent the blood from clotting on the striatum surface. A small amount of uncured Kwik-sil was applied to the striatum surface and a custom-made cannula window (1.8 mm in outer diameter, 1.4 mm in depth, Supplementary Fig. 2A) was carefully lowered into the brain. The outer ring of the cannula window was then fixed to the skull with instant adhesive (LOCTITE 495, Henkel) and dental cement. The inside of the cannula was then filled with Kwik-sil as a protector. After surgery, mice were allowed to recover with full access to water for at least 7 days before behavior training was resumed.

### Two-photon calcium imaging

Calcium imaging was performed using a custom built two-photon microscope (https://wiki.janelia.org/wiki/display/shareddesigns/MIMMS). The entire microscope was enclosed in a custom-designed, double-walled sound-attenuating box to eliminate potential influence from ambient noise of the two-photon imaging system (internal noise level < 30 dB SPL with the two-photon imaging system running). The noise produced by the resonant scanner were attenuated to < 30 dB SPL by an optical window sealing the output opening of the resonant scanner module. GCaMP6s was excited using a Ti-Sapphire laser (Chameleon UltraII, Coherent) tuned to 920 nm. Images were acquired using a 16X 0.8 NA objective (Nikon), and the GCaMP6s fluorescence was isolated using a bandpass filter (525/50, Semrock), and detected using GaAsP photomultiplier tubes (10770PB-40, Hamamatsu).

Horizontal scanning was accomplished using a resonant galvanometer (Thorlabs; 16 kHz line rate, bidirectional). The average power for imaging was ∼70 mW, measured at the entrance pupil of the objective. The field of view was ∼500×500 μm (at zoom 2×) or ∼350×350 μm (at zoom 3×) imaged at frame rate of ∼28 Hz for 512×512 pixels (Bi-direction) or 256×512 pixels (Single-direction). The acquisition system was controlled by ScanImage (https://scanimage.org) [6]. For each mouse, the optical axis was adjusted (40∼50 degree from vertical) to be perpendicular to the imaging window. Different fields of view in the same mouse were imaged on different sessions. The imaging was acquired continuously within each session, while the ScanImage system received a tag signal from the PX-Behavior System at the initiation of each trial (after ITI).

### Surgical procedures for fiber photometry and optogenetics

To prepare animals for fiber photometry recordings or optogenetic manipulations, mice were anaesthetized with isoflurane (1∼2%), and a small craniotomy (∼400 μm in diameter) was made vertically above the target area in each hemisphere.

For fiber photometry recording from dSPNs axons in SNr, 300 nL AAV2/9-hSyn-FLEX- GCaMP6s was slowly injected in bilateral TS (AP: -1.46, ML: 3.25, DV: -3.4) of D1-Cre mice, and fiber optics (NA 0.37, 200 μm core, ferrule O.D., 1.25mm) were inserted into the bilateral SNr (AP: -3.5, ML: 1.8, DV: 4.3) (**Fig. 4**, Supplementary Fig. 14). For fiber photometry recording from the PV neurons, 300 nL AAV2/9-hSyn-FLEX-GCaMP6s was injected into bilateral TS of PV-Cre mice, and fiber optics (NA 0.37, 200 μm core, ferrule O.D. 1.25mm) were inserted ∼200 μm above the virus injection sites (**Fig. 7A**, Supplementary Fig. 14B).

For optogenetic manipulations of specific types of striatal neurons, 400 nL AAV2/8-CAG-FLEX- ChR2(H134R)-mCherry (for activation, Supplementary Fig. 14C, 2.58×10^13^ V.G./mL) or 400 nL AAV2/9-Ef1α-DIO-eNpHR3.0-eYFP (for inactivation, Supplementary Fig. 14D, 1.50×10^13^ V.G./mL) virus was slowly injected into TS (AP: -1.46, ML: 3.25, DV: -3.4) in D1-Cre, D2-Cre or PV-Cre mice (**Fig. 5**, **Fig. 6G-6I**, **Fig. 7D-7F**). For iSPN inactivation by stGtACR2, AAV2/9- hSyn-SIO-stGtACR2-FusionRed (**Fig. 6G-6I**) was injected into TS in D2-Cre mice. For non-selective inactivation of striatal neurons, 400 nL AAV2/9-CAG-ArchT-GFP (3.50×10^12^ V.G./mL) virus was injected to TS or to an anterior sub-region of dorsal stratum (AP: -0.02, ML: 2.0, DV: - 3.4) in wildtype C57BL/6J mice (Supplementary Fig. 14E). For control animals, AAV2/9-CAG-eGFP-2A (2.40×10^13^ V.G./mL) virus was injected into TS (Supplementary Fig. 14F). About 30 min after virus injection, fiber optics for optogenetic manipulation (NA 0.37, 200 μm core, ceramic ferrule OD 1.25mm) were implanted ∼200 μm above the virus injection sites.

After injection of virus and implantation the fiber optics, silicon adhesive (Kwik-sil, WPI) was used to cover the craniotomy. A custom-designed titanium headplate was attached to the skull with dental cement for head fixation. After surgery, mice were allowed to recover at least 7 days before water restriction and behavioral training.

### Fiber photometry recording

The fiber photometry system was custom built based on previous method [7]. The excitation light was from a 470-nm LED (Cree, Inc, XQEBLU-SB-0000- 00000000Y01-ND) and a 410-nm LED (Xuming, 3W, 410-420 nm). The intensity of the 470-nm light was set to 10-15 μW measured at the distal end of the patch cord (calibrated based on the transmittance of the fiber optics implanted in the brain, to keep the intensity at 10 μW around the fiber optic tips). For each recording site, the intensity of 410-nm light was adjusted to keep the recorded emission fluorescence level excited by 410-nm light close to that excited by 470-nm light from the brain tissue.

The lights from 470-nm LED and 410-nm LED independently passing through aspheric condenser lenses (Thorlabs, ACL25416), were filtered through respective bandpass filters (FB470-10, and FB410-10, Thorlabs), and were then combined by a dichroic mirror (DMLP425R, Thorlabs). The combined excitation lights then sequentially passed through a dichroic mirror (Chroma, T525/50dcrb) and an objective (Olympus, UPLFLN 20X NA0.5), and then were coupled into a fiber optic cable (NA 0.37, 200 μm core) that affixed to the fiber optics implanted in mouse brain. A small fraction of 470-nm and 410-nm lights reflected by the first dichroic was collected by a condenser lens (Thorlabs, ACL25416) and received by a photodiode (Thorlabs, PAD100A2), the real-time light intensity of which was used as a feedback signal for custom-designed closed-loop circuits to stabilize the intensity of 470-nm and 410-nm lights throughout the recording session (**Fig. 4A**).

The emitted fluorescent signals were collected by the same light path and focused onto a CCD (QImaging, R1 retina). The 470-nm LED, the 410-nm LED and the CCD were controlled by a NI I/O device (NI, USB-6341) and software written in LabVIEW. The NI device also receives the triggers from the digital outputs of PX-Behavior System to tag the behavior-related time.

Consecutive camera frames were captured at 40 Hz using alternating 470-nm and 410-nm excitation lights (20 Hz sampling rate for both 470 nm and 410 nm channels). The excitation pulse duration of 470-nm LED or 410-nm LED in each frame was 11 ms for an exposure time of 10 ms, beginning at 0.5 ms prior to the onset of the exposure time of CCD and ending at 0.5 ms after the offset of the exposure time of CCD. The signals from CCD were recorded by ImageJ based software Micro-Manager (Version 2.0 beta), with the ‘binning’ at 16x16, and the ‘Exposure’ time of 10ms. The fluorescent signals excited by the 470-nm and 410-nm were extracted and analyzed subsequently using MATLAB scripts.

### Optogenetic manipulations

For cell-type specific activation by ChR2, a 473-nm diode-pumped solid state laser (BL473T8-100FC, Shanghai Laser & Optics Century Co., Ltd.) with the output power adjusted to ∼2 mW at the distal end of patch cable was used to activate ChR2-expressing neurons. For cell-type specific inactivation by NpHR3.0, a 593-nm diode-pumped solid state laser (YL593T6-100FC, Shanghai Laser & Optics Century Co., Ltd.) with output power of ∼20 mW was used to inactivate NpHR-expressing neurons. For iSPN inactivation by stGtACR2, a 462-nm blue diode laser (BLM462TA-300FC, Shanghai Laser & Optics Century Co., Ltd.) with an output power of 2 mW was used to inactivate stGtACR2-expressing iSPNs. For iSPN activation by ChrimsonR, a 635-nm red diode laser (RLM635TA-100FC, Shanghai Laser & Optics Century Co., Ltd.) with an output power of 5 mW at the distal end of patch cable was used to activate ChrimsonR-expressing iSPNs. For inactivation experiments using ArchT and control group expressing GFP (Supplementary Fig. 1D), a 532-nm diode-pumped solid state laser (GL532TA-100FC, Shanghai Laser & Optics Century Co., Ltd.) with an output power of 20 mW at the distal end of patch cable was used.

In all optogenetic stimulation sessions, photostimulation was applied to a proportion of trials randomly interleaved with control trials (no photostimulation) to allow for within-subject and within-session comparison. Since in most sessions there were less trials with intermediate frequencies (∼30%) and more trials with the end frequencies (∼70%), to obtain comparable number of trials with photostimulation for intermediate and end frequencies, photostimulation was applied to ∼50% of the intermediate-frequency trials, and ∼15% of end-frequency trials.

### Histology

After the completion of experiments, mice were deeply anesthetized and perfused with saline and 4% PFA (or 10% formalin), and brains were removed and post-fixed overnight. Brains were then incubated with PBS containing 30% sucrose overnight. Frozen sections (50 µm, using Leica CM1950 cryostats) containing viral injection sites, cannula window location and optic fiber tracks were collected and stained with DAPI to visualize cell nuclei. Histology images were taken using Olympus VS120® Virtual Microscopy (Supplementary Fig. 14). For verification of cell-type specific virus expression in dSPNs and iSPNs, 200 nL AAV2/8-CAG-FLEX-eGFP-WPRE-pA (1.25×1013 V.G./mL, TaiTool Bioscience) was injected into the TS of D1-Cre or D2-Cre mice. One month after the surgery, the animals were perfused and the brains were sectioned sagittally to examine the projection pattern of neurons expressing GFP (Supplementary Fig. 2B-C).

### Two-photon imaging data analysis

All imaging data analysis was performed by custom-written MATLAB (Mathworks) routines. Within each session, the brain motion artifacts were corrected by registering each frame to a target image using a cross-correlation-based registration algorithm (discrete Fourier transformation (DFT) algorithm) [8]. The target image was the averaged projection of several visually identified frames with minimal brain motion artifacts. Individual neurons as regions of interest (ROI) were visually identified using the averaged projection of images from each trial. The average fluorescence values of all pixels within each ROI across frames in one session were used as the raw fluorescence data (F_raw_) of each neuron. The slow fluorescence change was removed by subtracting the 8^th^ percentile value of F_raw_ within nearby 500 frames for each frame (forward 250 frames and backward 250 frames, ∼17.5 s). The mode of all these 8^th^ percentile values within one session was then added back to restore the fluorescence levels of each neuron, resulting in the corrected fluorescence data (F). The calcium signal (ΔF/F) was calculated as:

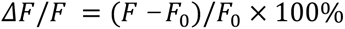

Where F_0_ is the baseline fluorescence defined as the middle value of the peak histogram of F (bin size, 500). Estimation of spike rate using calcium signal deconvolution was based on the Fast Nonnegative Deconvolution algorithm [9].

To quantify single neuronal selectivity differentiating trial types with ipsilateral and contralateral choices, we used an ideal observer decoding based on ROC analysis [10]. By comparing the distribution of the neuronal responses in contralateral choice trials and those in ipsilateral trials (with respect to the imaged hemisphere), the area under ROC curve (auROC) value for choice quantifies how well a neuron could differentiate between contralateral choice and ipsilateral choice. We then calculated the selectivity index (SI) [11] to indicate the relative preference for contralateral vs ipsilateral trials:

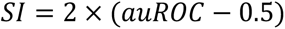

where value 1 indicates the strongest preference for contralateral choices and value -1 for ipsilateral choices.

The significance level of SI was defined by a permutation analysis, where we shuffled the trial label (contralateral or ipsilateral choice) for 1000 times and each time we calculated an SI. Values greater than 99.5th percentile or smaller than 0.5th percentile of the shuffled results are considered as p<0.01. The SI values were obtained using a generalized ROC algorithm[12]. To calculate the SI, mean ΔF/F of each trial during 100 ms preceding answer time was used in **Fig. 2A-2B**, and mean inferred spike rate during 100 ms preceding answer time was used in **Fig. 2H-2I**. Neurons with significant contralateral-preference in correct trials and neurons with significant ipsilateral-preference in correct trials were included in **Fig. 2A-2B, 2H-2I**. In Supplementary Fig. 3, contralateral-preferring neurons: with significant positive SIs both in correct and error trials; ipsilateral-preferring neurons: with significant negative SIs both in correct and error trials; sensory category preference: with opposite significant preference in correct and error trials.

To quantify the temporal correlation between simultaneously imaged neurons, we calculated the Pearson’s correlation coefficient (CC) between the time series of calcium signals from pairs of neurons (**Fig. 3**). Averaged CCs were reported as the average across neuronal pairs (**Fig.3C**). In **Fig. 3E-3G** and Supplementary Fig. 4, all neurons in the imaging field are used to calculate CCs. In **Fig. 3H-3K**, only neurons with significant choice-selectivity were included. The cross-correlation between neuronal pairs were obtained by compute the Pearson’s correlation coefficients between the ΔF/F traces from pairs of neurons with different time lags, ranging from -10 s to +10 s, which were then plotted as a function of the time lag (**Fig. 3C** and **3D**). We do not exclude the possibility that mixing of optical signals from neuropils, which cannot be optically isolated, could potentially contribute to the level of correlation [13]. However, for dSPNs or iSPNs, the neuropils are all from the genetically defined cell types, thus the cell-type specific difference in the correlation coefficients still reflects the difference in the level of local population coordination between dSPNs and iSPNs.

### Fiber photometry data analysis

The 410-nm reference trace was scaled to best fit the 470-nm signals using least-squares regression [14]. Then the motion-corrected 470-nm signal (F_470_) was obtained by subtracting the scaled 410-nm reference trace from the 470-nm signal and then adding the mean value of scaled 410-nm trace. ΔF/F signal was calculated by subtracting the mode of F_470_ and then dividing by the mode value of F_470_.

To compare signals across recording sessions, we used the z-score of calcium signals, which is the ΔF/F subtracted by the mean and divided by the standard deviation of the data in the session. The peak value of z-score within 1 s after answer time was used for comparison (**Fig. 4C, 7C**). To quantify the encoding of choice direction (Contra. vs Ipsi. choice) in the neuronal activity measured with fiber photometry, mean ΔF/F of each trial during 1 s after answer time was used to calculate the selectivity index (SI) in each recording session. Statistical significance was determined with a permutation test at the significance level of α = 0.05 (**Fig. 4D**).

### Optogenetics data analysis

For analyzing the behavioral effects of optogenetic stimulation, ‘miss’ trials were excluded. To quantify the behavioral effects of optogenetic stimulation, we used the two-way repeated-measures ANOVA with function ‘fitrm’ in Matlab to test the effect across tone frequencies, and use LSD post hoc tests to compare the effects at each frequency.

### Statistical Analysis

All statistical analysis was performed using MATLAB 2017a (MathWorks). All data are shown as mean ± s.e.m. and shaded regions surrounding line-plots indicate s.e.m. unless mentioned otherwise. The comparison significance was tested using Kolmogorov– Smirnov test, one-sample proportion test, paired two-sided Wilcoxon signed rank test, two-way repeated-measures ANOVA and post hoc Fisher’s LSD multiple comparison tests. *P < 0.05, **P < 0.01, ***P < 0.001. For fiber photometry recording from SNr, two sessions were excluded due to the inaccuracy of fiber optics tip location. For activation of iSPNs, one session was excluded due to inconsistent sound frequency setting with other sessions.

### Neural network model

To examine the potential computational implementation for how concurrent but asymmetrical dSPN and iSPN activity could result in unambiguous action selection following decisions, we constructed a computational model of the cortico-basal ganglia circuitry on the population level, which is similar to the computational model of dopamine- biasing action selection through basal ganglia in the previous study [15]. In the model, the decision outcomes are transferred from cortex to dSPNs, iSPNs and striatal PV neurons via canonical cortico-striatal projections [16]. The activity of the dSPNs and iSPNs populations converge to the ipsilateral substantia nigra pars reticulata (SNr) populations through the direct and the indirect pathway respectively. The left and right SNr populations mutually inhibit each other, and the dominant SNr population determines the motor output by controlling the final motor output either through brainstem circuits or motor cortices. The network model composed of neural populations in both left and right hemispheres with symmetrical connectivity, as shown in **Fig. 8A**.

In the model, a perceptual decision G is produced in the cortex according to a simple decision rule (Equation 1). Where *sign*(*q*) = 1 if *q* ≥ 0, *sign*(*q*) = −1 if *q* < 0. Note that the tone of stimulus *q* ∈ [−1,1] is in the unit of octaves. *θ*(*q*) is the probability of wrong decision due to perceptual uncertainty, where *σ* = 2.24 is the degree of uncertainty in the perception (Equation 2). The decision outcome *d* = ±1 encodes left or right choice. The cortical neurons in each hemispace are organized into two types of populations that are selective to contralateral or ipsilateral choices. The firing rate of cortical population *CTX_jk_* is shown in Equation 3, where *j* ∈ {*L*, *R*} representing the left or right hemisphere, *k* ∈ {*contra*, *ipsi*} is the selectivity for left (*s* = −1) or right (*s* = 1) action, *s* = −1 corresponds to *f^CTX_Rcontra_^* and *f^CTX_Lipsi_^*, *s* = 1 corresponds to *f^CTX_Lcontra_^* and *f^CTX_Ripsi_^* and *d* is the decision outcome given by Equation 1. *I* = 20 is the peak firing rate of cortical neurons. *f_b_* = 0 is reference frequency. As the frequency *q* changes, the firing activity of cortical neural population switches between low and high activity states in a sigmoid form, with the switching slope controlled by ζ = 0.01. Cortical neurons keep self-sustained activity states for a period of time without external input [17].

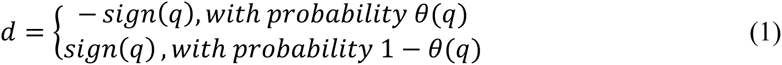

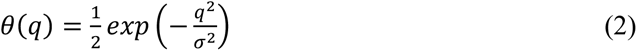

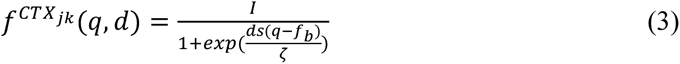

SPNs in the striatum receive currents both externally and internally (Equation 4), where 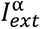 and 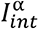 are the external current and internal current to population *α*, from the cortex and from striatal sub-populations respectively (**Fig. 8A**). SPNs are divided into 8 sub-populations based on different response preferences found in our imaging results and from the two hemispheres, *α* = *D_ijk_*, with *i* ∈ {1,2} representing cell types (dSPNs and iSPNs). The cortical input to population *α* is shown in Equation 5, where *f^CTXjk^* is the activity of the cortical population with selectivity *s* (Equation 3). 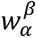 is the connection strength from population *β* to population *α* (Supplementary table 1). *V_α_* is the membrane potential. For the excitatory connection from *CTX_jk_*, the reverse potential *E_CTXjk→α_* = 0. The internal input current to population α is shown in Equation 6, where *β* is a striatal population including α and PV, and *f^β^* is its firing activity. Reverse potential parameters are set according to precious study [18]. For inhibitory connections from striatal population *β*, the reverse potential was *E_β_*_→α_= −65*mV*. For the connection from iSPN to SNr, *E_D2jk→SNrj_*= 0 *mV*.

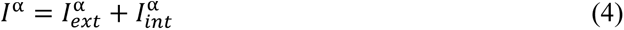

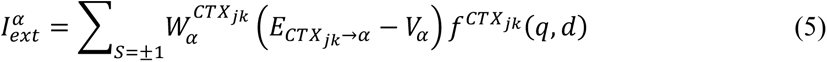

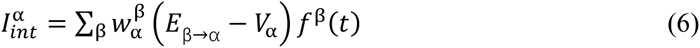

Take *D_1,LContra_* as an example of SPN sub-population to explain its specific input current shown in Equation 4-6,

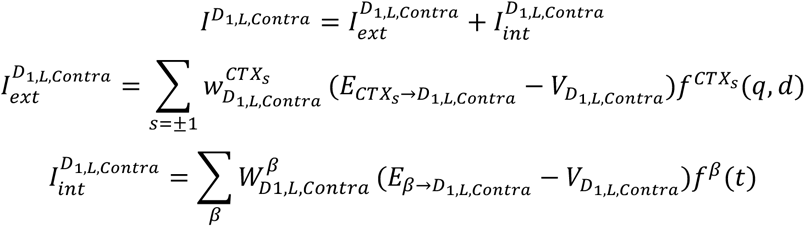

PV neurons only receive cortical inputs from ipsilateral cortex, 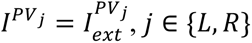, take PV neurons in the left hemisphere as an example,

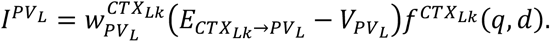

SNr neurons receive external current from SPNs and internal current as their inputs. Take SNr neurons in the left hemisphere as an example,

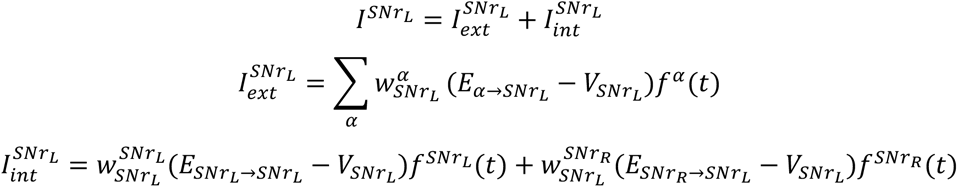

The membrane potential of a striatal population α is described as shown in Equation 7. *E_α_* = −55 *mV* is the reversal potential. ɛ is a noise term. The firing frequency of a striatal population α is shown in Equation 8. The transfer function *g* is linear above the discharge threshold θ = −40 *mV* and has an exponential growth trend below the discharge threshold (Equation 9), where *k* = 1 represents spiny projection neurons, and Z = 5 represents interneurons in the striatum.

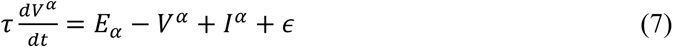

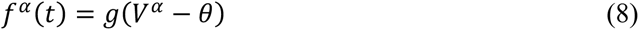

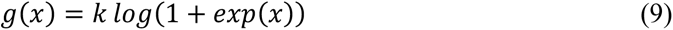

The motor choice is made according to the SNr activity after 100 ms when the network activity becomes stabilized. The rule for motor choice is defined in Equation 10.

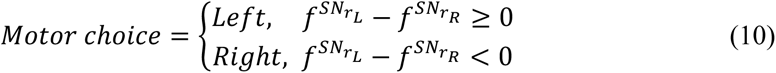

The connection weights between the populations in the network are shown in Supplementary table 1. The weights are assumed to form through pervious learning processes. Populations are labeled according to the convention introduced in previous sections. For example, *CTX_−_*_1_ is the cortical population selective to left choice, and *D_1Rcontra_* is the dSPN population connected to the right SNr and is selective for contralateral choice. For simplicity, we omitted some indices of the labels if they are not concerned. For example, *D_1L_* includes populations *D_1Lipsi_* and *D*_1*Lcontra*_. The dSPNs and iSPNs in the striatum inhibit each other, and the weights are shown as 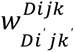.

The optogenetic manipulation of a striatal population *α* is modelled by an additional current *I^stim^* to the membrane potential dynamics (Equation 11), where *I^stim^* is the regulation current, taking positive or negative values for optogenetic activation or inactivation, respectively. The parameters of optogenetic regulation are shown in Supplementary table 2.

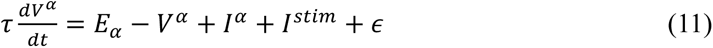

To simulate the condition that dSPN and iSPN populations show identical activities, we assigned the connection weights from cortical populations to dSPN populations to the connection weights from cortical populations to the corresponding iSPN populations,

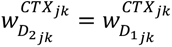

We performed the network model stimulation using the same set of parameters under the standard control conditions (Supplementary table 1), as well as under the perturbed conditions caused by optogenetic activation or inactivation (Supplementary table 2). Under each condition, 300 independent trials were simulated, and the probability and the standard deviation of choosing right choice was calculated. Psychometric functions were fitted by a 4-parameter sigmoidal curve same as in the experimental data. Compared with the standard control situation, under the perturbed condition of optogenetic activation/inactivation of striatal PV neurons, dSPNs, and iSPNs, the model reproduced similar biased action selection behavior as observed under the corresponding conditions in experiments (**Fig. 8D-8H**). The psychometric functions of action selection of the model qualitatively captured the deviation from that of the control condition after perturbation as shown in experiments.

---

**Supplementary Fig. 1.**
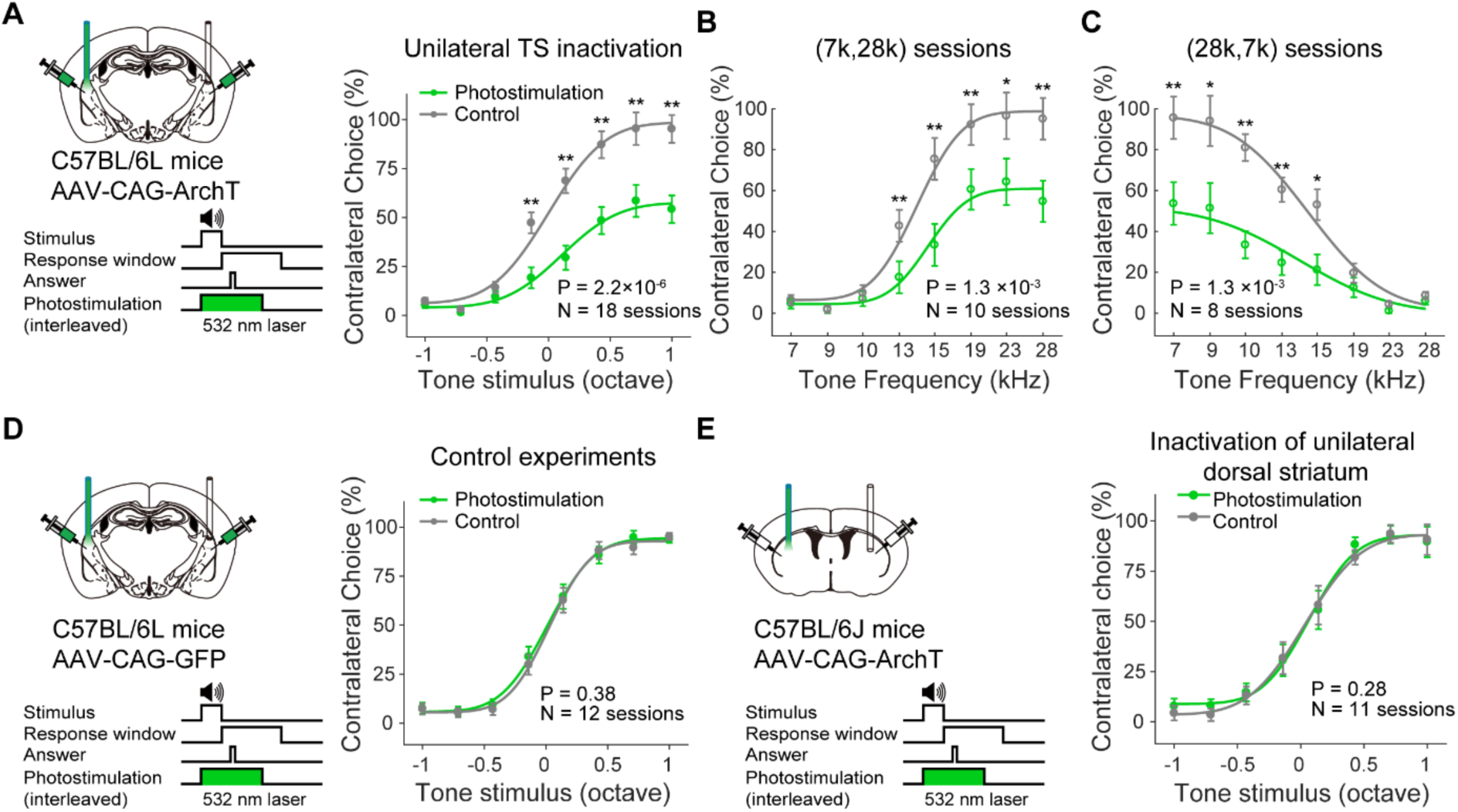
Unilateral optogenetic inactivation of the TS and the anterior sub- region of the dorsal striatum. (A) AAV-CAG-ArchT was used to inactivate the TS during the auditory decision task. Summarized psychometric functions from all sessions (n = 18 sessions from 9 mice). (B-C) Psychometric functions from two groups of mice trained with reversed stimulus-choice association resepctively. n = 10 sessions from 5 mice in (B). n = 8 sessions from 4 mice in (C). (D) Similar as in (A), in control group injected with AAV-CAG-GFP in the TS, n = 12 sessions from 3 mice. (E) Similar as in (A) for optogenetic inactivation of the anterior sub- region of the dorsal striatum, injected with AAV-CAG-ArchT, n = 11 sessions from 6 mice. P values shown are for effects of photostimulation from two-way repeated-measures ANOVA. * p < 0.05, ** p < 0.01, Fisher’s LSD post hoc tests.

**Supplementary Fig. 2.**
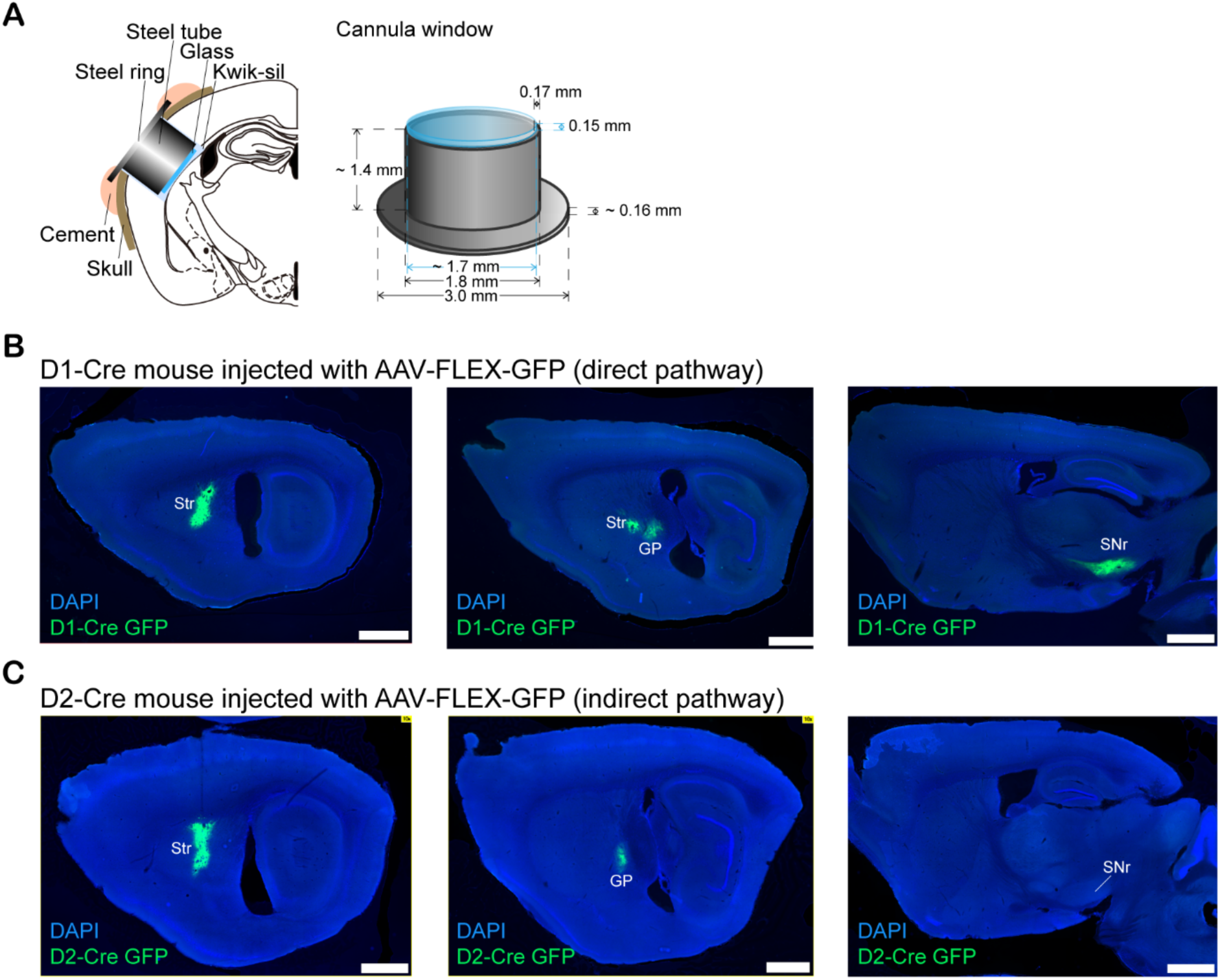
Cannular window for chronic in vivo two-photon calcium imaging from the posterior tail of the striatum (TS) and histology verifying the cell type specific virus labeling of SPNs. (A) Schematic showing chronically implanted cannular window for *in vivo* two-photon imaging of the TS. (B) Sagittal section from an example D1-Cre mouse, showing Cre-dependent viral expression (GFP) in the TS and downstream axonal targets (GP and SNr) of labeled dSPNs. (C) Sagittal section from an example D2-Cre mouse, showing Cre-dependent viral expression (GFP) in the TS and downstream axonal targets (GP only) of labeled neurons. Scale bars, 1 mm.

**Supplementary Fig. 3.**
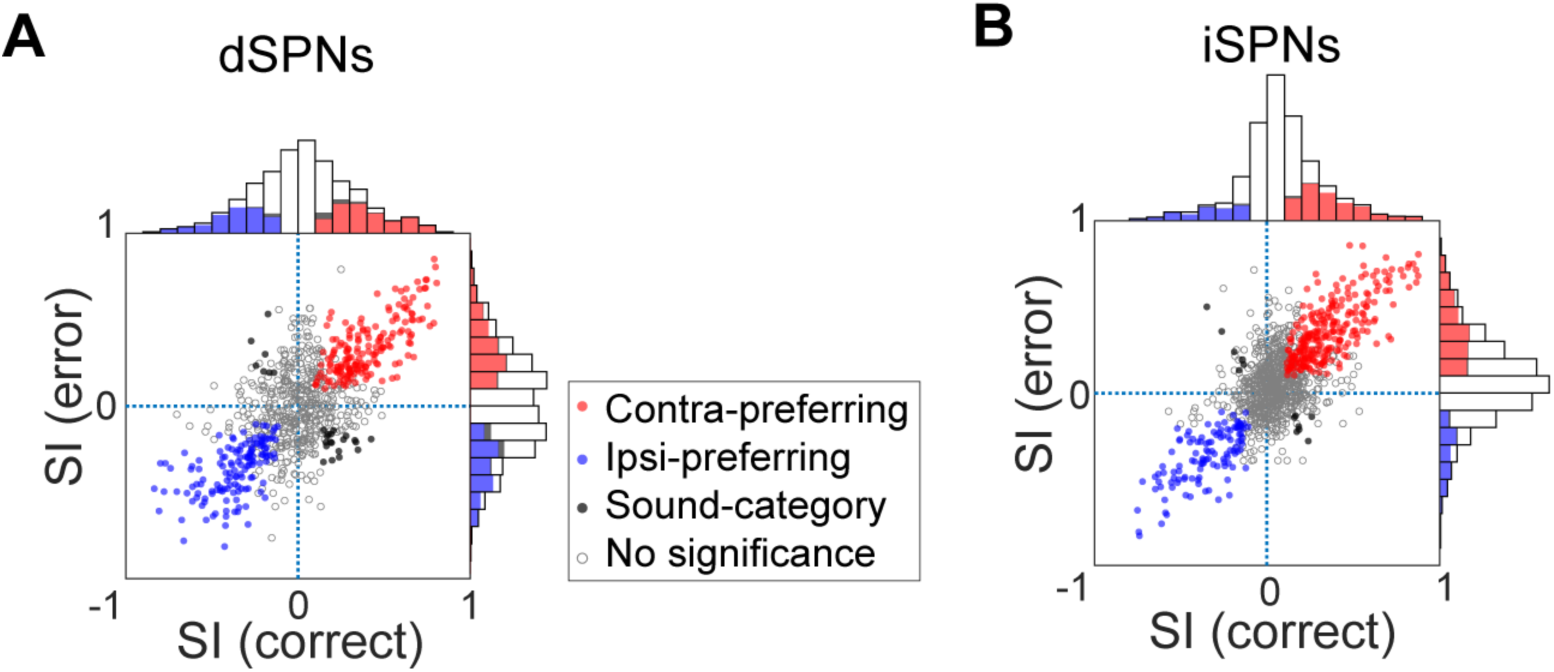
dSPNs and iSPNs encode choice direction. (A) Index of selectivity (SI) for choices calculated using averaged calcium signals in a 100 ms window before answer time. SIs from correct trials were plotted against SIs from error trials for individual dSPNs, n = 811 neurons from 8 imaging fields of 4 mice. (B) Similar as in (A) for iSPNs, n = 1152 neurons from 11 imaging fields of 6 mice. Red, neurons showing significant contralateral preference in both correct and error trials; Blue, neurons showing significant ipsilateral preference in both correct and error trials; Black, sensory category selective neurons; Grey, below significance level in either correct or error trials.

**Supplementary Fig. 4.**
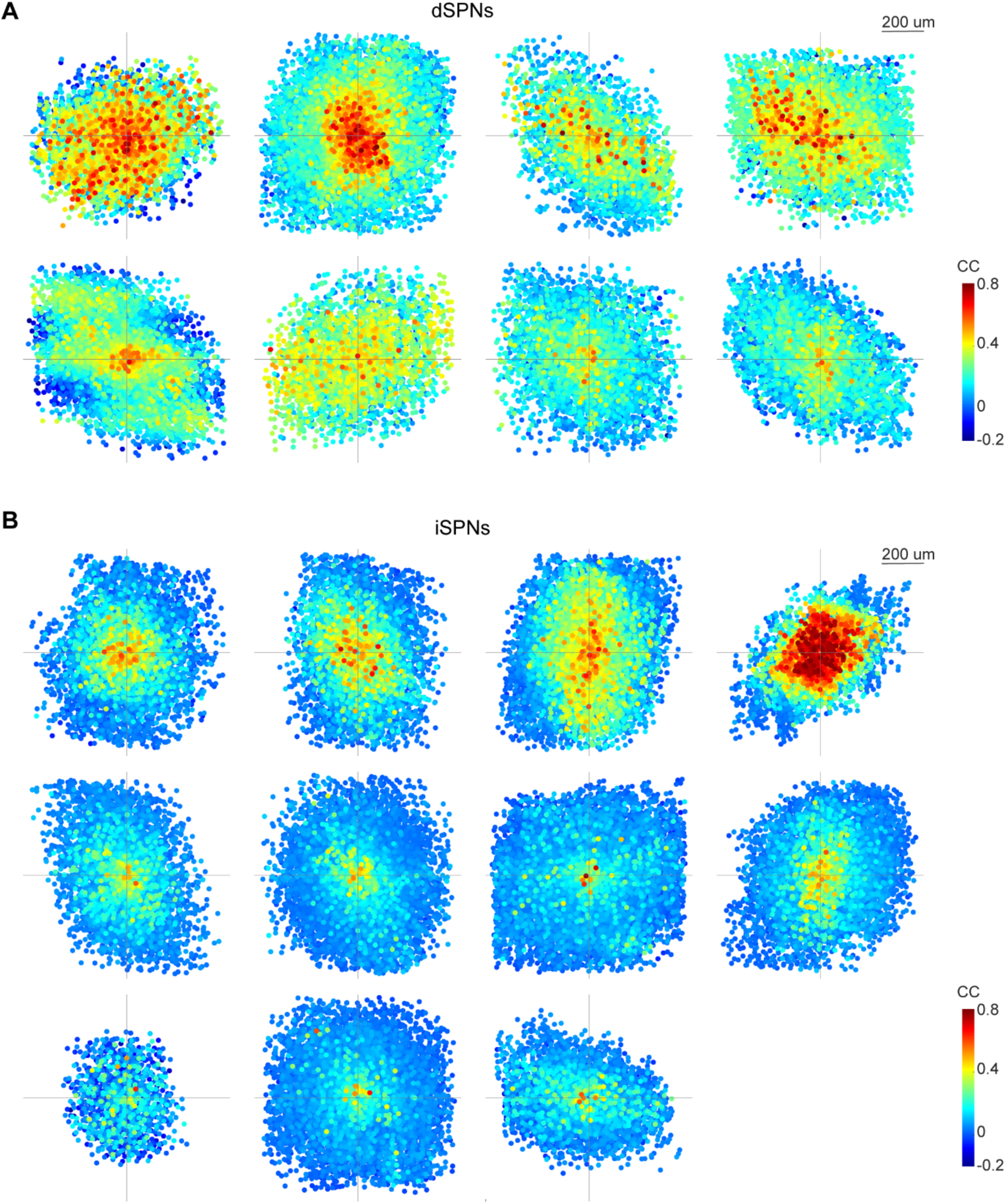
Spatiotemporal correlation of dSPNs and iSPNs for individual imaging fields. (A) Spatiotemporal correlation of dSPNs from 8 imaging fields of 4 D1-Cre mice, similar as in Fig. 3E. (B) Spatiotemporal correlation of iSPNs from 11 imaging fields of 6 D2-Cre mice.

**Supplementary Fig. 5.**
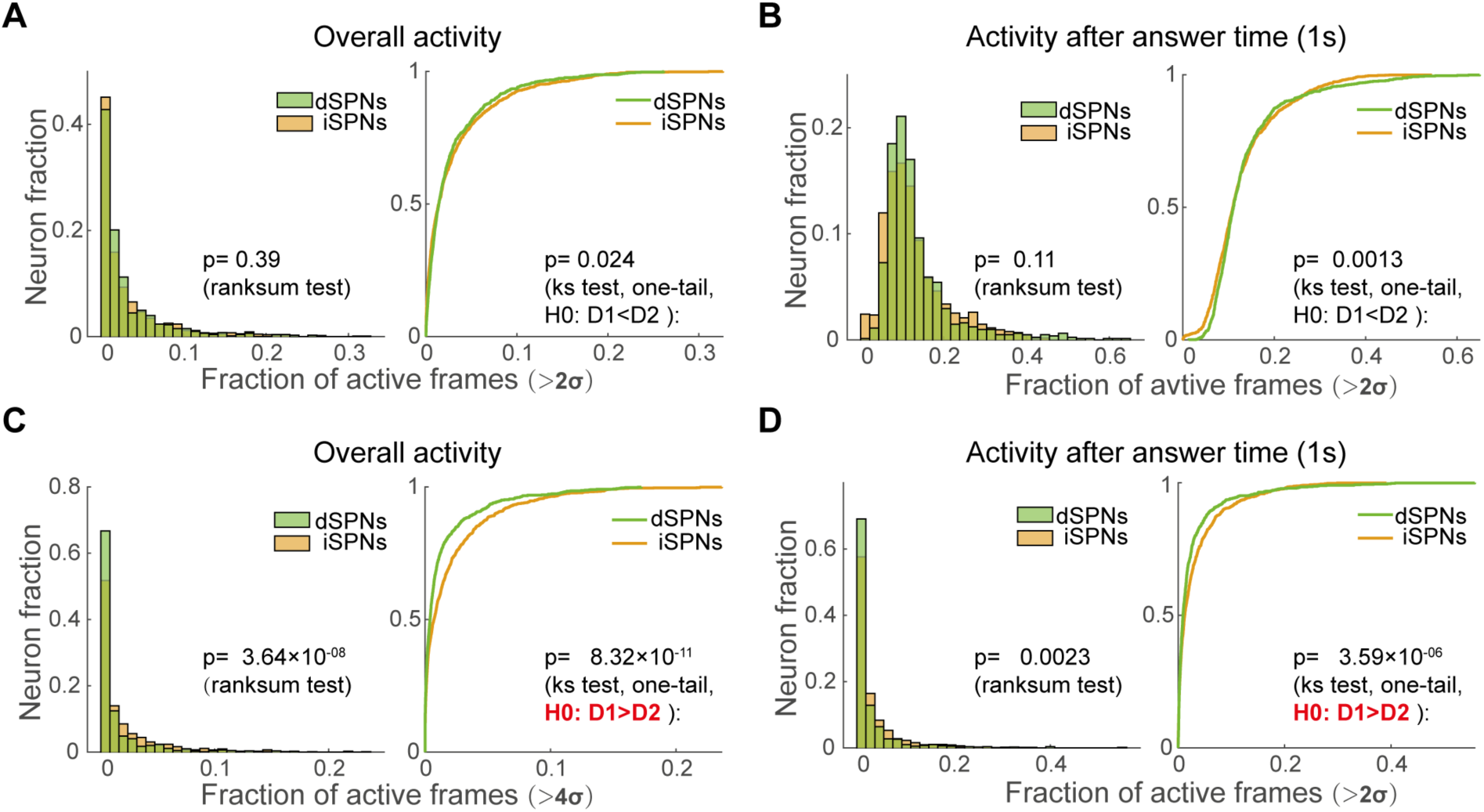
The activity level of dSPNs and iSPNs during sensory motor decision-making. (A) Comparison of activity level (fraction of acive frame) for dSPNs (green) and iSPNs (orange). Left, the distribution of activity level; Right, the CDF curve. The criterion of active frame was that consecutive over 5 frames (about 175 ms) were above 2σ. The data used for analysis are consecutive calcium signal from whole behavior session. (B) Similar as in (A), but the data used for analysi are within the task-related period (1 second from mice answer time). (C-D) Similar as in (A-B), but criterion of active frame was that consecutive over 5 frames were above 4σ.

**Supplementary Fig. 6.**
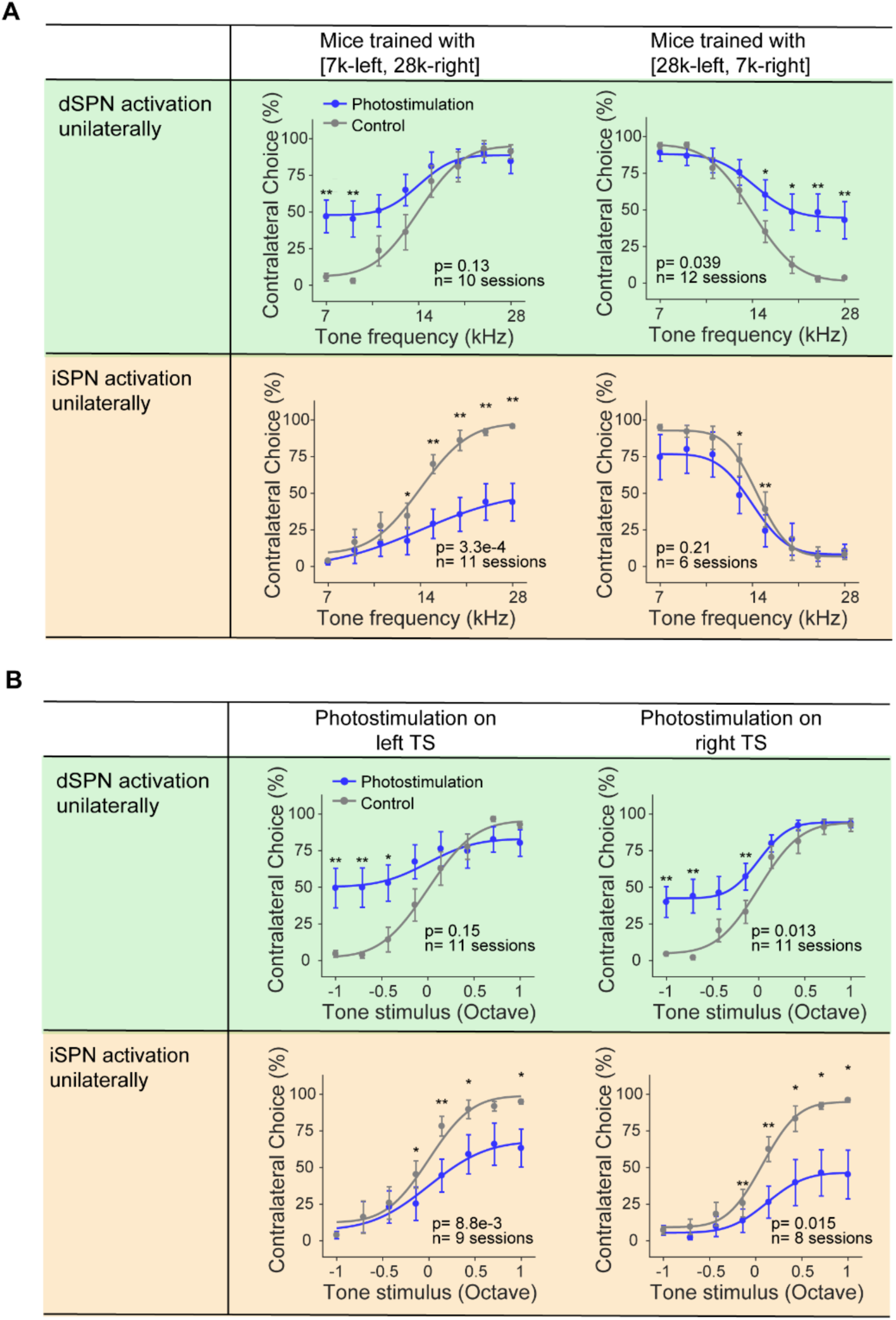
The effects of optogenetic activation with different sound-choice association and in left/right hemisphere respectively. (A) Psychometric functions from the two groups of mice trained with reversed stimulus-choice associated were summarized separately, showing percentages of choices contralateral to the photostimulated hemisphere as a function of the tone frequency values. Top two are from dSPN activation, and bottom two are from iSPN activation. (B) Summarized psychometric functions from control trials (grey) and trials in the left TS and in the right TS, showing percentages of choices contralateral to the photostimulated hemisphere as a function of the tone stimulus. The tone stimuli were arranged such that the positive values indicate tones associated with contralateral choices and negative values associated with ipsilateral choices. Top two are from dSPN activation, and bottom two are from iSPN activation.

**Supplementary Fig. 7.**
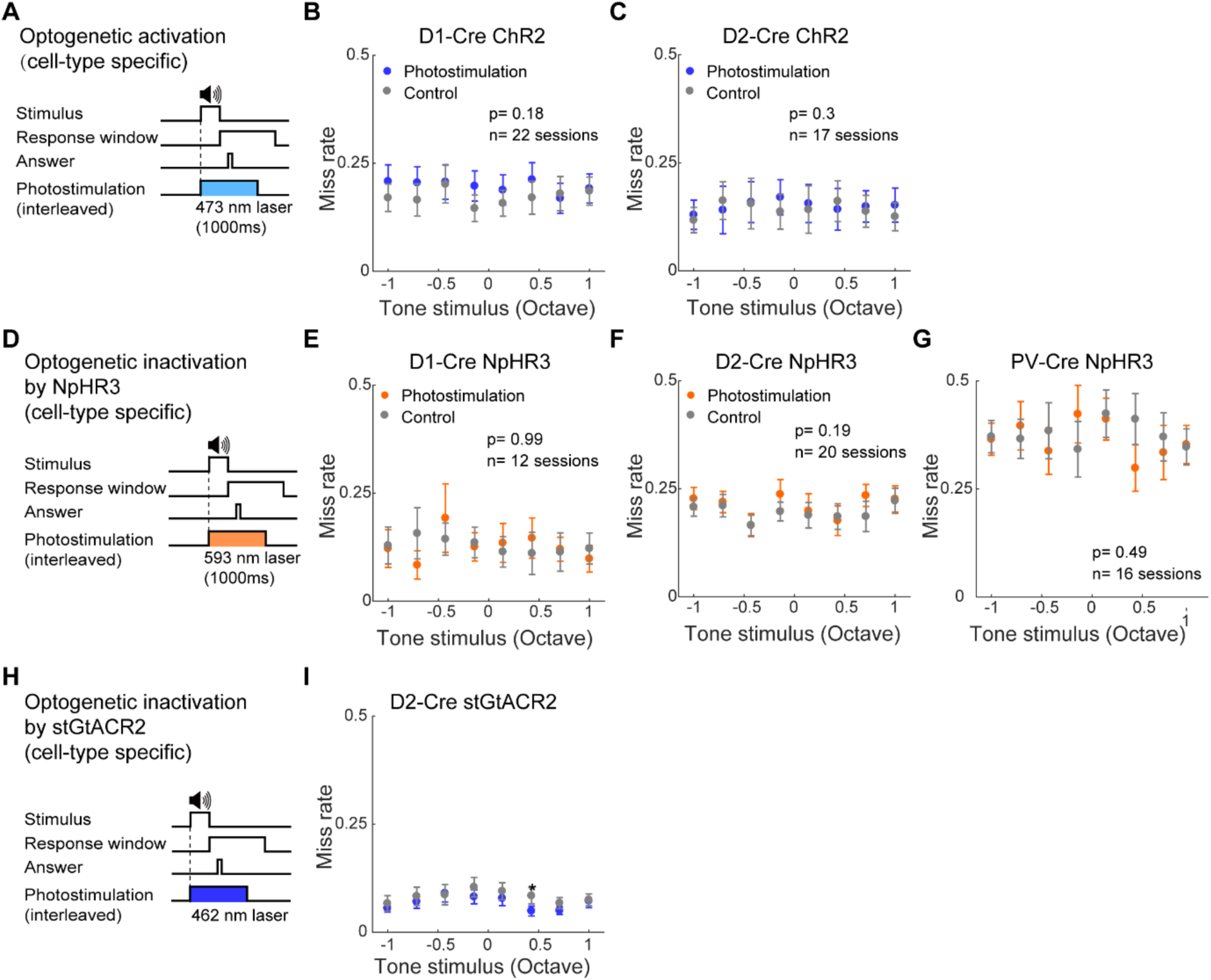
The effects of optogenetic manipulations on miss rate. (A) Schematic showing trial structure with optogenetic activation. (B-C) Miss rate as a function of tone stimulus in trials with or without optogenetic activation of dSPNs (B) or activation of iSPNs (C). (D) Schematic showing trial structure with optogenetic inactivation using eNpHR3.0. (E-G) Similar as in (B) for optogenetic inactivation of dSPNs (E), inactivation of iSPNs (F), inactivation of PV interneurons (G). (H) Schematic showing trial structure with optogenetic inactivation of iSPNs using stGtACR2. (I) Similar as in (F) for optogenetic inactivation of iSPNs using stGtACR2. P values shown are for effects of photostimulation from two-way repeated-measures ANOVA.

**Supplementary Fig. 8.**
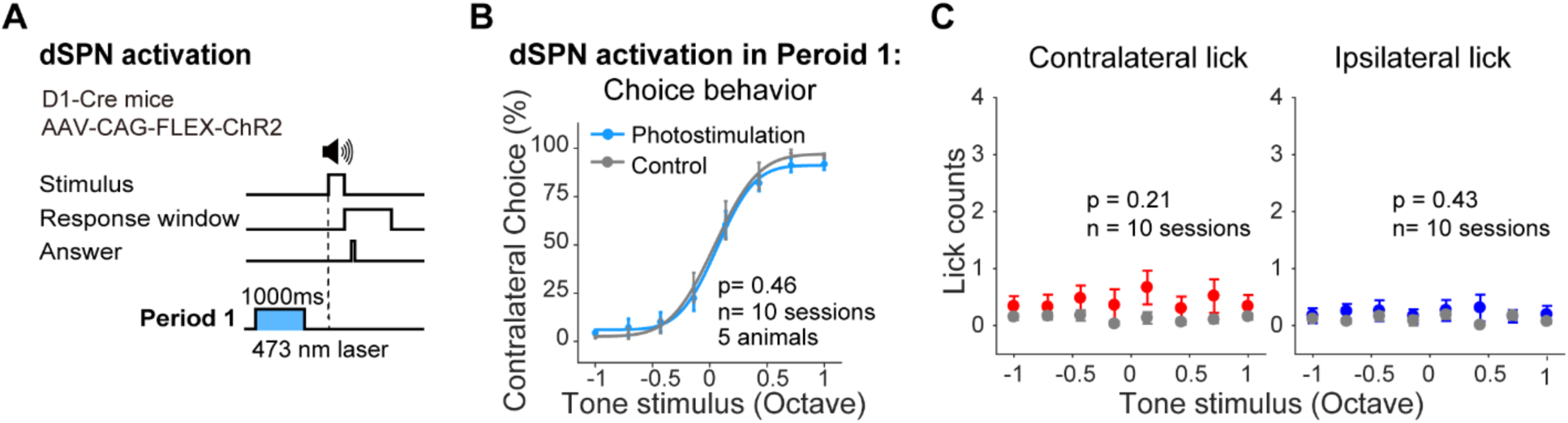
The effect of dSPN activation out of sound period. (A) Schematic showing trial structure with optogenetic activation of dSPNs before sound onset (during inter-trial-interval). (B) Summarized psychometric functions from control trials (grey) and trials with dSPN activation (blue) in period 1 as shown in (A). (C) Number of licks within the period of photostimulation as a function of tone stimulus from trials with or without photostimulation. Only correct trials were included. Left, contralateral licking. Right, ipsilateral licking.

**Supplementary Fig. 9.**
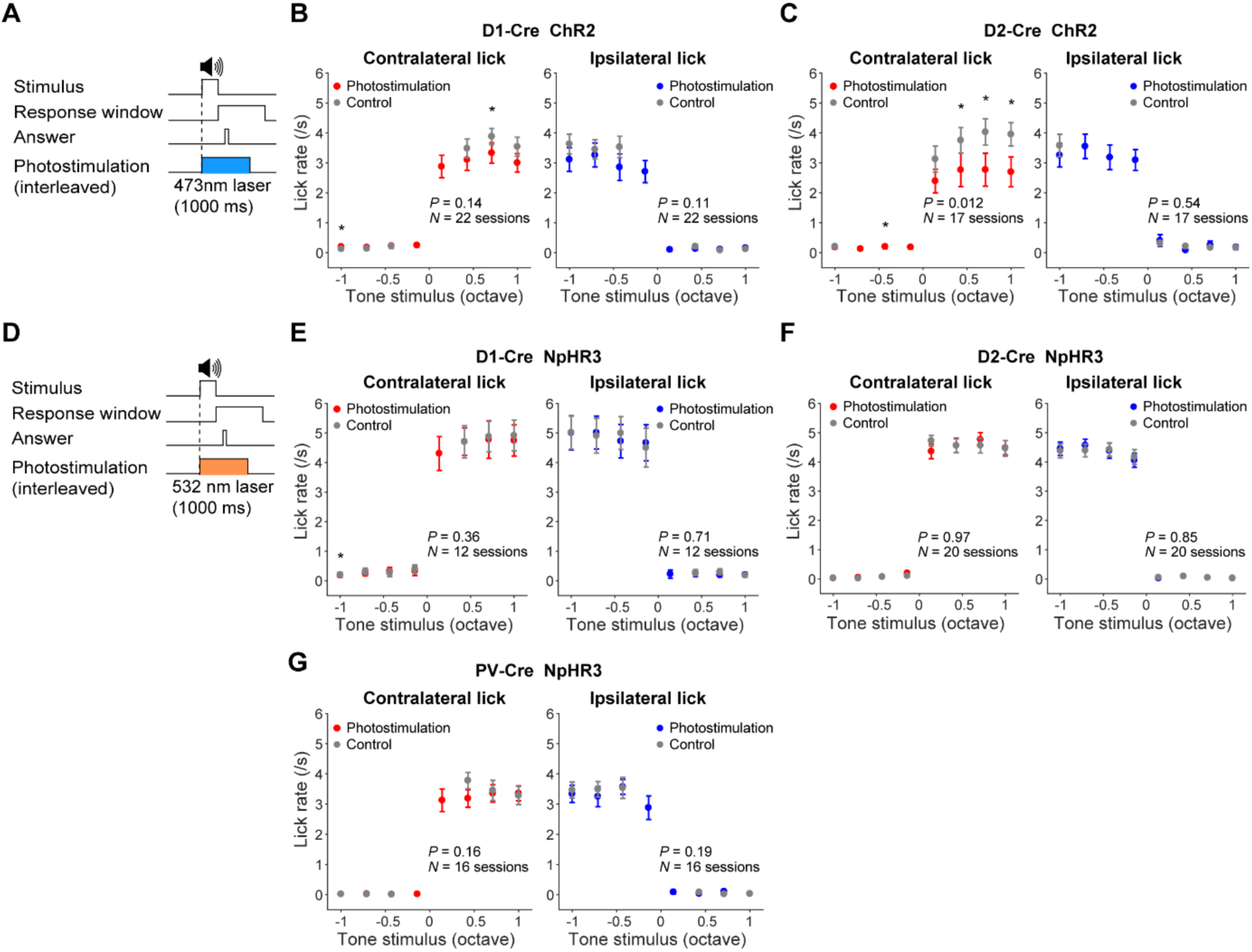
The effects of optogenetic manipulations on lick rate. (A) Schematic showing trial structure with optogenetic activation of dSPNs or iSPNs. (B) Number of licks within the period of photostimulation (activivation of dSPNs) as a function of tone stimulus from trials with or without photostimulation. Only correct trials were included. Left, contralateral licking. Right, ipsilateral licking. (C) Similar as in (B) for activation of iSPNs. (D) Schematic showing trial structure with optogenetic inactivation of dSPNs, iSPNs or PV neurons using NpHR3. (E) Similar as in (B) for optogenetic inactivation of dSPNs. (F) Similar as in (B) for optogenetic inactivation of iSPNs. (G) Similar as in (B) for optogenetic inactivation of PV neurons. P values shown are for effects of photostimulation from two-way repeated-measures ANOVA.

**Supplementary Fig. 10.**
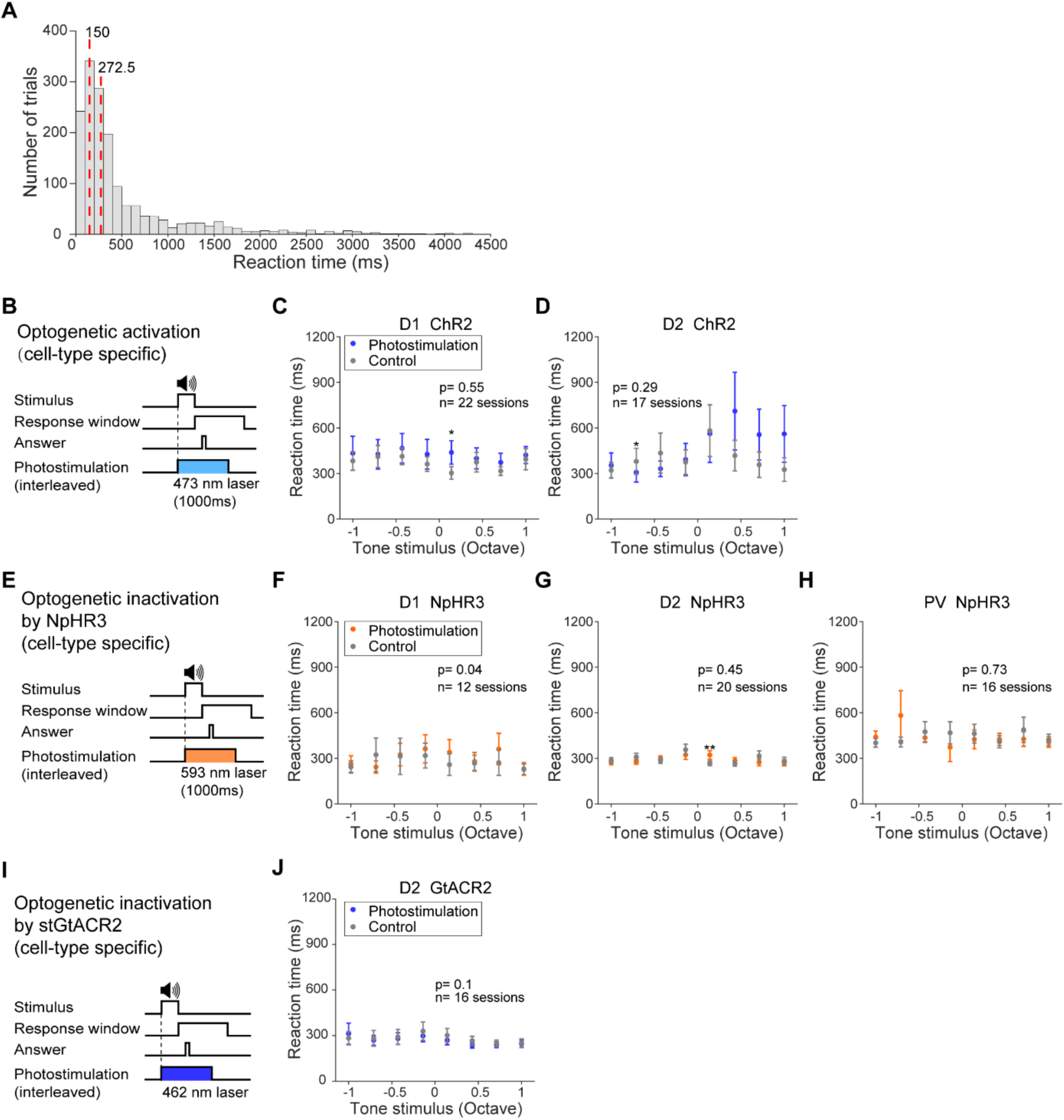
The effects of optogenetic manipulations on mice’s reaction time. (A) Distribution of reaction time in control trials for D2-Cre mice as shown in Figure 5D. Vertical red lines indicate the first quarter (150 ms) and the median (272.5 ms). (B) Schematic showing trial structure with optogenetic activation. (C) Reaction time as a function of tone stimulus in trials with or without optogenetic activation of dSPNs. (D) As in (C) for iSPN activation. (E) Schematic showing trial structure with optogenetic inactivation with eNpHR3.0. (F-H) Similar as in (C) for dSPN inactivation (F), iSPN inactivation (G), and PV interneuron inactivation (H). (I) Schematic showing trial structure with optogenetic inactivation with stGtACR2. (J) Similar as in (C) for iSPN inactivation with stGtACR2. P values shown are for effects of photostimulation from two-way repeated-measures ANOVA. * p < 0.05, Fisher’s LSD post hoc tests.

**Supplementary Fig. 11.**
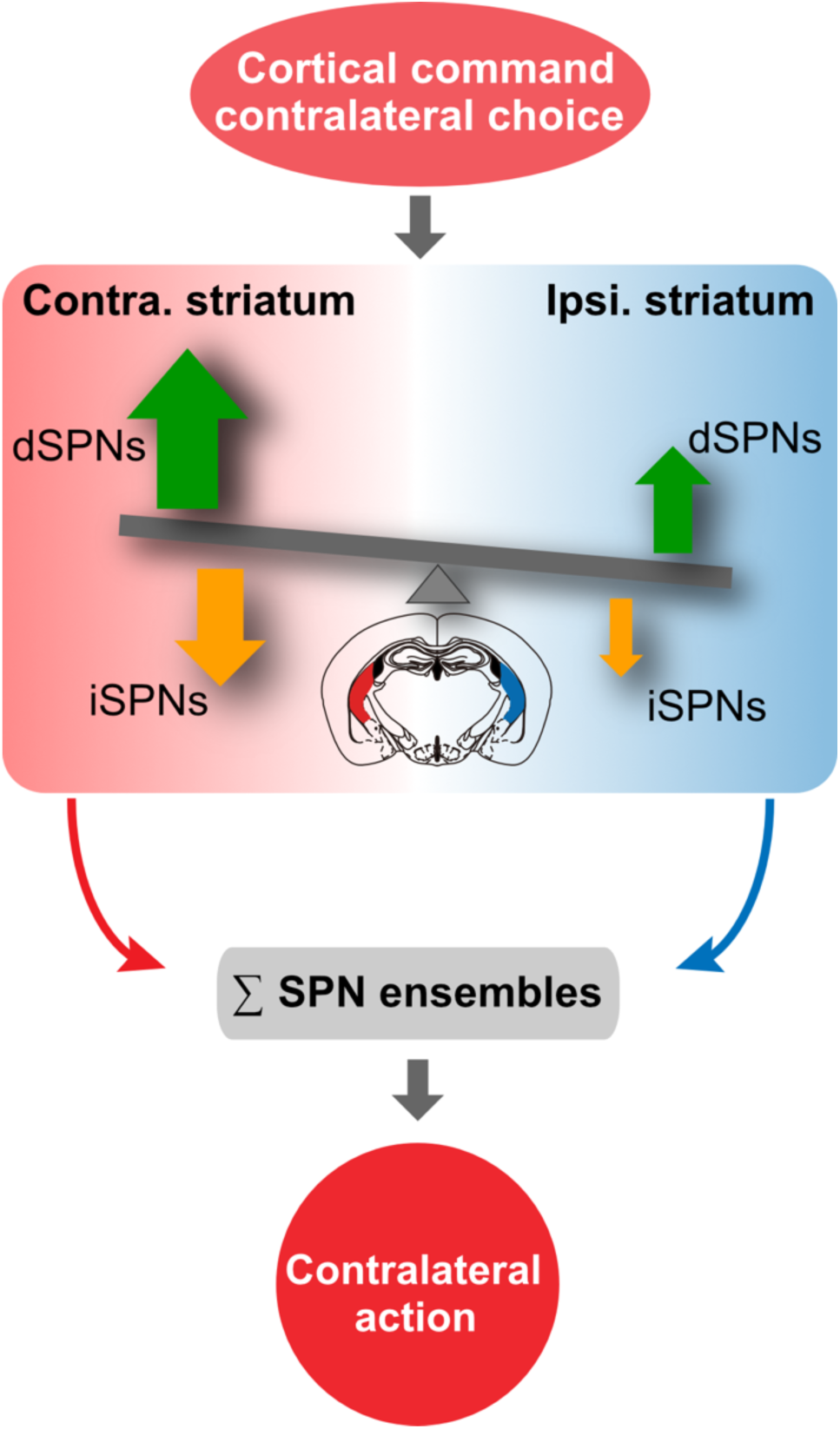
Schematic showing a conceptual model with balanced functional ensembles from both dSPNs and iSPNs that control action selection. The striatum receives dynamic inputs from cortex and thalamus that could be balanced or biased between competing choices. Within dSPNs or iSPNs there are subgroups of neurons preferring different choices and each can bidirectly regulate action selection. At ensemble level, the dSPNs in the tail of the striatum (TS) show predominantly stronger coding strength for the contralateral choice, which may produce unambiguous action selection following population pooling.

**Supplementary table 1.**
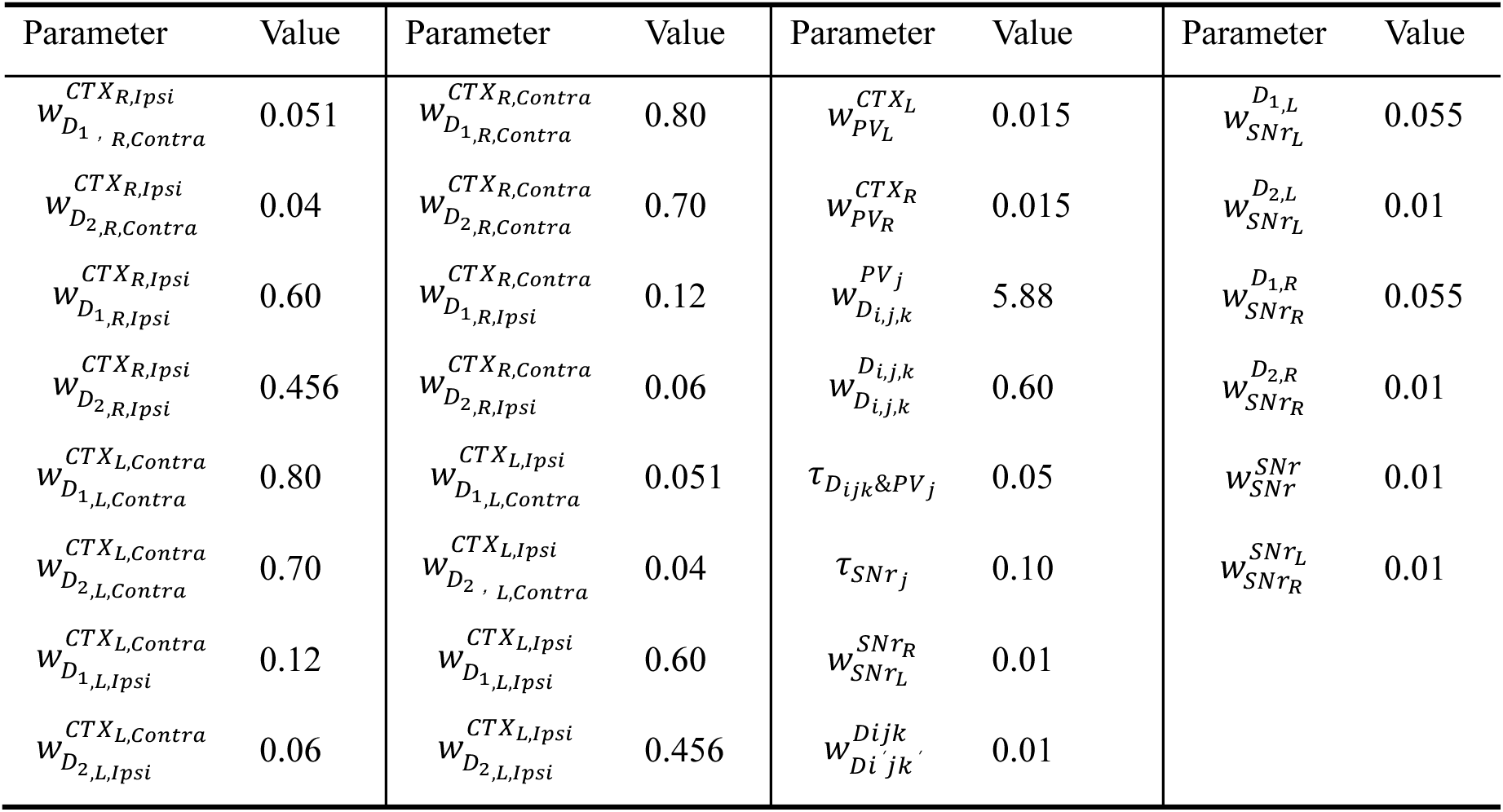
Parameters for the network model, related to Fig. 8.

**Supplementary table 2.** Optogenetic perturbation current of left hemispace (μA), related to Fig. 8.

| Type | D1 activation | D2 activation | D1 inactivation | D2 inactivation | PV inactivation |
| --- | --- | --- | --- | --- | --- |
| $I^{stim}$ | 10.5±2.5 | 10.5±2.5 | -2.5±1.0 | -2.5±1.0 | -2.5±1.0 |

**Supplementary movie 1: Example video of two-photon imaging during behavioral task.** Video clip showing continuous two-photon imaging from the TS of the left hemisphere during the head-fixed mouse performing the behavioral task. The OLED board on the right showing trial information and performance outcome.

**Supplementary movie 2: Example video of optogenetic stimulation during behavioral task.** Video clip showing optogenetic activation of dSPNs in the right TS during the mouse performing the behavioral task. The OLED board on the right showing trial information and performance outcome. Note that the photostimulation in the third trials shown was accompanied by a wrong choice of the animal.

## Notes

### Competing Interest Statement

The authors have declared no competing interest.

### Summary of Updates

Main text updated for improved clarity; Supplemental files updated.

